# Microbial evolution reshapes soil carbon feedbacks to climate change

**DOI:** 10.1101/641399

**Authors:** Elsa Abs, Scott R. Saleska, Regis Ferriere

## Abstract

Microbial decomposition of soil organic matter is a key component of the global carbon cycle. As Earth’s climate changes, the response of microbes and microbial enzymes to rising temperatures will largely determine the soil carbon feedback to atmospheric CO2. However, while increasing attention focuses on physiological and ecological mechanisms of microbial responses, the role of evolutionary adaptation has been little studied. To address this gap, we developed an ecosystem-evolutionary model of a soil microbe-enzyme system under warming. Constraining the model with observations from five contrasting sites reveals evolutionary aggravation of soil carbon loss to be the most likely outcome; however, temperature-dependent increases in mortality could cause an evolutionary buffering effect instead. We generally predict a strong latitudinal pattern, from small evolutionary effects at low latitude to large evolutionary effects at high latitudes. Accounting for evolutionary mechanisms will likely be critical for improving projections of Earth system responses to climate change.

---

Microorganisms are key drivers of global biogeochemical cycles^1^. In terrestrial ecosystems, soil microbes decompose organic matter, returning carbon to the atmosphere as carbon dioxide (CO_2_)^2^. *In vitro* and *in situ* experiments suggest that changes in microbial decomposition with warming are an important feedback to climate^3–5^. Soil microbial populations may respond to increasing temperature through physiological mechanisms such as individual metabolic adjustment^6,7^ and ecological mechanisms such as shifts in population abundance or community composition^8,9^. Given the short generation time, large population sizes and standing genetic variation of many microbial organisms, evolutionary adaptive responses of microbial populations to warming are also likely^10,11^. However, how microbial evolutionary adaptation may contribute to carbon-climate feedbacks is unknown^12^.

Key to microbial decomposition of soil organic matter is the production by microbes of extracellular enzymes (exoenzymes), that diffuse locally in the soil and bind to soil organic matter compounds^13^. Because the fitness cost of exoenzyme production^14^ (reduced allocation to growth, Fig. 1a) is paid by individual microbes whereas fitness benefits (larger resource pool) are received by microbial collectives^15^, we expect genetic variation in exoenzyme production^16^ to be under strong selection^15,17^. Our objective is to evaluate how exoenzyme production responds to selection under environmental warming, and how the evolutionary response of exoenzyme production impacts the response of soil organic carbon stock (SOC).

**Figure 1.**
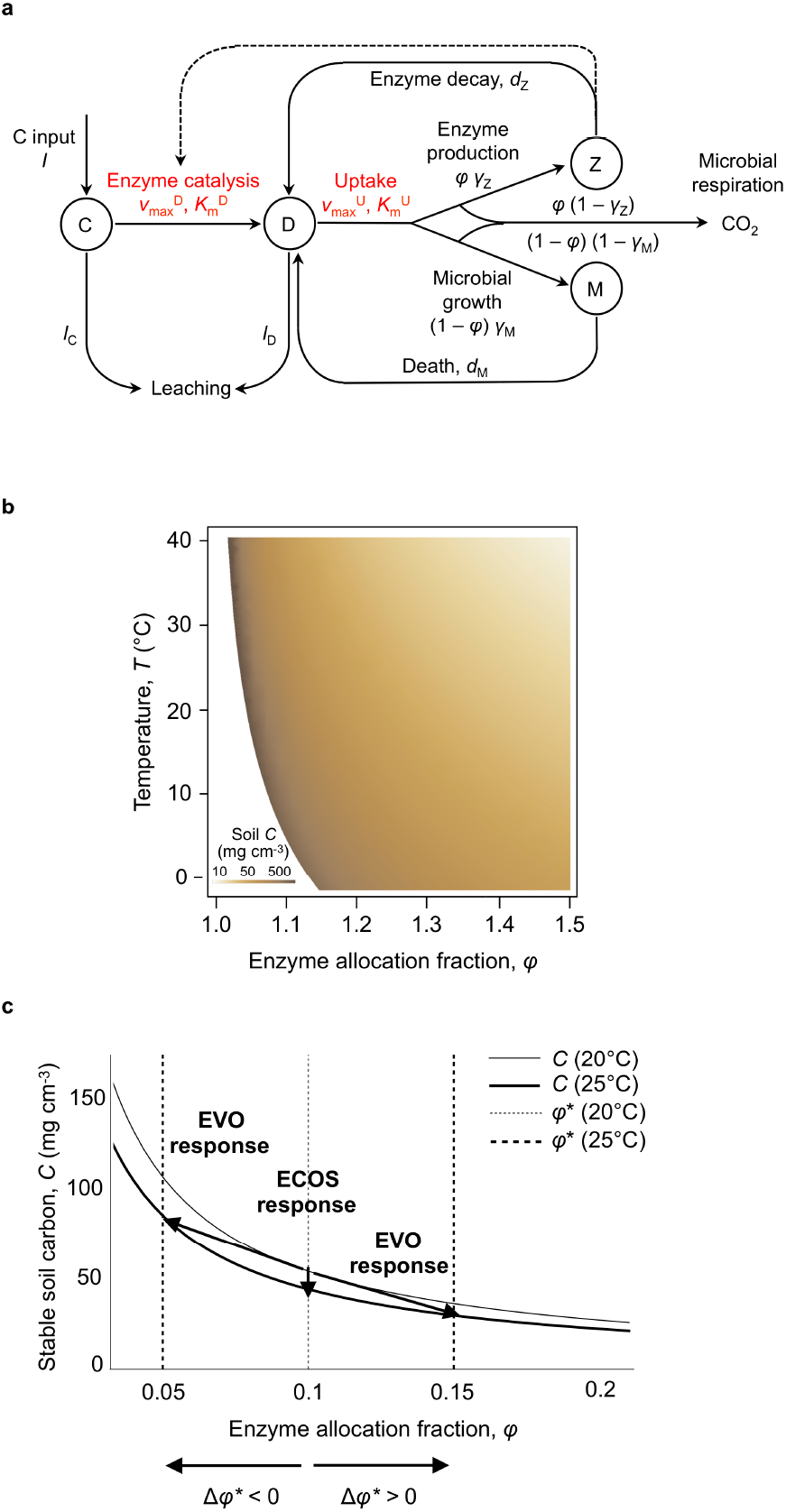
Effect of temperature and enzyme allocation fraction on SOC ecosystem equilibrium. **a**, Structure of the microbial-enzyme ecosystem model (see Methods for details): SOC stock is the balance of plant input, *I*, and loss by exoenzyme-mediated degradation to DOC (*D*), which in turn is allocated between (fraction *φ*) production of enzymes (*Z*) and (fraction 1— *φ*) growth of microbial biomass (*M*). **b**, Effect of temperature and enzyme allocation fraction, *φ*, on SOC equilibrium, *C*, in the baseline scenario of temperature dependence. **c**, Response of SOC ecosystem equilibrium, *C*, to a 5°C increase in temperature (from 20 °C to 25 °C) as a function of enzyme allocation fraction, *φ*. Parameters are set to their default values (Supplementary Table 1), except *I* = 5 10^−3^, 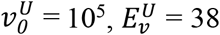 and *c*_0_ = 1.17.

To this end, we develop and analyze a novel ecosystem-evolutionary model, starting with an ecosystem model of microbe-enzyme decomposition first proposed by Allison *et al.* (2010)^18^ (Fig. 1a), and then modified to take microbial evolutionary adaptation into account. Amongst ecosystem models of soil microbial decomposition (reviewed in Abs and Ferriere (2019)), Allison *et al.^18^*’s model is the simplest of the *CDMZ* type, where *C* denotes a single pool of soil organic carbon, D, dissolved organic carbon, M, the biomass of the microbial community, and *Z*, a single pool of exoenzymes. The focal microbial trait is the fraction of assimilated carbon allocated to exoenzyme production^19–21^, or ‘exoenzyme allocation fraction’, whose mean value across microbial strains or species is denoted by *φ*. The balance of assimilated carbon, 1 – φ, is allocated to microbial growth. In the absence of evolution, the enzyme allocation fraction, *φ*, represents the mean of an unchanging distribution of enzyme allocation behaviors across a community of microbes. With natural selection acting on the trait’s genetic variation across the community, *φ* evolves as the microbial community adapts to the environmental conditions, e.g. mean soil temperature. Our ecosystem-evolutionary model predicts the evolutionarily stable value, *φ*\*, as a function of temperature, and how the response of *φ*\* to warming affects the decomposition rate and SOC stock (*C* at equilibrium).

By comparing the response of the SOC stock that our ecosystem-evolutionary model predicts (EVO response) to the response predicted by the ecosystem *CDMZ* model in the absence of evolution, assuming *φ* to be a fixed parameter (ECOS response), we can evaluate the contribution of microbial evolutionary adaptation (EVO effect) to the direction and magnitude of the SOC stock response to climate warming (Fig. 1b, c). To illustrate how EVO effects may vary in real ecosystems, we use available data^22^ on the decomposition kinetic parameters in five sites of increasing latitude and decreasing mean annual temperature. We evaluate ECOS and EVO responses for each site, and compare them within and among sites. Our analysis identifies parameters and temperature dependencies that critically influence the strength of evolutionary effects. We discuss how these evolutionary effects relate to previous consideration of ‘adaptation’ in microbial community responses to climate warming^18,23,24^, and how evolutionary adaptation may interact with ecological responses such as species sorting and community shifts^25–27^. We conclude by highlighting how our results could inform future empirical work.

## Model overview

We use the microbe-enzyme model of litter decomposition first introduced in ref. 18^18^ to describe the ecosystem dynamics of soil organic carbon (*C*), dissolved organic carbon (*D*), microbial biomass (*M*), and extracellular enzyme abundance (*Z*), given litter input, leaching rates, and soil temperature (Fig. 1a and Supplementary Figure 1). The effect of temperature is mediated by enzymes kinetics, with exoenzymes driving the decomposition rate, and intra-cellular enzymes involved in resource uptake and microbial biomass synthesis. The model parameters (microbial life history parameters, thermal dependencies, enzyme parameters, carbon inputs) were constrained by experimental and observational data^18,20,22^ (see Abs and Ferriere 2019 for a review). Within these parameter ranges, the model outputs are consistent with target empirical values (target *C* of the order of 100 mg cm^−3^, *M* about 2% of C, *Z* about 1% of *M*, and limiting *D* close to zero). Zhang *et al.^28^* provided some more direct validation by successfully fitting a *CDMZ* model to time series of field measurements of soil respiration from a specific ecosystem (semiarid savannah subject to episodic rainfall pulses).

In general, as temperature increases, the baseline ecosystem model without evolution predicts^18^ a decline in equilibrium SOC due to faster enzyme kinetics, a positive feedback to warming. We call this the ECOS (for ECOSystem-only) response of the system. To investigate the effect of evolutionary adaptation on decomposition, we include microbial evolution in the ecosystem model. Our focus is on soil bacteria (as opposed to fungi), which typically have large population size and short generation time. DOC uptaken by individual cells is allocated to exoenzyme production (fraction *φ*) or microbial biomass, with enzyme production efficiency denoted by *γ_Z_* and microbial growth efficiency (MGE) denoted by *γ_M_*. We assume that microbes may vary individually in their exoenzyme allocation fraction and that some of this variation has a genetic basis. We then derive the selection gradient on the enzyme allocation fraction *φ* and compute the evolutionarily stable value, *φ*\*. Changing temperature alters the selection gradient, hence *φ*\*. Knowing how *φ*\* changes as temperature increases, we can evaluate how the ecosystem equilibrium changes from both the direct effect of warming on enzyme kinetics, and the indirect effect mediated by microbial evolutionary adaptation to warming (Fig. 1b, c).

At the molecular and cellular level, the effect of warming on microbial decomposition is mediated by the temperature sensitivity of intra- and extra-cellular enzymatic activity^22,29,30^. In our baseline ‘kinetics-only’ scenario of temperature-dependent decomposition, we assume that microbial uptake parameters (maximum uptake rate and half-saturation constant) and exoenzyme kinetics parameters (maximum decomposition rate and half-saturation constant) increase with temperature^5,31^ in a logistic manner. We consider two additional scenarios for the influence of temperature on decomposition. In the microbial mortality scenario, the microbial death rate also increases with temperature^32^.

This could be due to a higher risk of predation or pathogenic infection at higher temperatures, or faster microbial senescence due to higher protein turnover^32^. In the microbial growth efficiency (MGE) scenario, MGE (the fraction of carbon allocated to growth that actually contributes to microbial biomass, as opposed to being released as CO2 via growth respiration) decreases with temperature^18,24^, which could be due to higher maintenance costs at higher temperature^33^.

## Results

### Evolution of the enzyme allocation fraction trait

The adapted exoenzyme allocation fraction, *φ*\*, corresponds to a maximum of microbial fitness relative to other trait values, and therefore depends on the parameters defining microbial net growth rate: MGE, maximum uptake rate, local competitive advantage to exoenzyme producers, or ‘competition asymmetry’, and mortality. From our model, *φ*\* at temperature *T* is derived (methods) as:

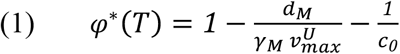

where *γ_M_* denotes the MGE; 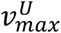, the maximum uptake rate; *d_M_*, the death rate, and *c*_0_, the degree of competition asymmetry. The latter quantifies the accessibility of the exoenzyme ‘public good’ to microbes; it measures the differential availability of enzymatically produced dissolved organic carbon (DOC) to different microbial strains. Competition asymmetry is shaped by diffusion of exoenzymes and DOC, and by microbial mobility, and is thus likely influenced by soil physical properties, such as texture or moisture. For simplicity, we assume that competition asymmetry is independent of temperature.

At a fixed temperature, the model predicts that microbes invest less in exoenzymes when exposed to hostile conditions, which may translate into high mortality (*d_M_*), low MGE (*γ_M_*), low maximal uptake rate 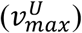, and/or low competitive advantage to producers (*c*_0_). Conversely, microbes are selected to allocate more resource into producing exoenzymes when more favourable growth conditions allow them to obtain a better ‘return on investment’^34^.

According to equation (1), the adaptive response of exoenzyme allocation to warming depends on how the MGE, maximum uptake, and mortality vary with temperature. We investigate how parameters influence the direction and sensitivity of the adaptive enzyme allocation fraction to temperature by calculating the derivative of *φ*\* with respect to *T*, for each scenario of temperature dependence. In the baseline ‘kinetics-only’ scenario (in which 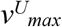 is the only variable in (1) that depends on temperature) the response of *φ*\* to temperature variation is given by

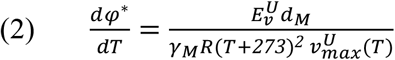

According to equation (2), increasing temperature creates more favorable growth conditions for microbes through higher resource uptake capacity (higher 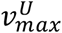), thus microbes evolve higher investment in exoenzyme production (positive *dφ*\*/d*T*) (Supplementary Figure 5a), resulting in lower equilibrium SOC (an evolutionary enhancement of the positive feedback to warming seen in the baseline ECOS response). Because *φ*\* is proportional to the product of the exponential of 1/T and to 1/*T*^2^, its sensitivity to temperature is strongest across low temperatures (equation (2), Supplementary Figure 5a). The adaptive response is generally predicted to be strongest (and the enhancement of the positive climate feedback greatest) when warming improves microbial growth potential (by increasing 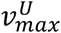) under initially hostile conditions (high *d_M_*, low *γ_M_*, low 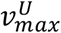 due to low intrinsic uptake rate 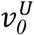 and/or high activation energy 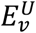, low initial temperature *T*_0_) (equation (2)).

In the temperature-dependent mortality scenario, both mortality and uptake respond exponentially to temperature. Microbes evolve a higher investment in exoenzyme production if the change in resource uptake capacity remains greater than the change in mortality rate; otherwise, microbes evolve a lower investment (Supplementary Note 3 and Supplementary Figure 5b-c). In the MGE-temperature dependent scenario, the response of *φ*\* to temperature variation is proportional to the inverse of an exponential function of temperature (through the uptake rate, 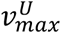) times a linear function of temperature^32^ (through MGE). As a consequence, at low initial temperature, the adaptive response of the allocation strategy to warming is mainly driven by the thermal dependence of resource uptake, and microbes adapt by increasing their resource investment in exoenzyme production. For ecosystems that are initially warmer, the microbial adaptive response to warming is mainly driven by the reduction of MGE, which leads to lower allocation to exoenzymes (Supplementary Note 3 and Supplementary Figure 5d).

There are two cases where the evolutionary enhancement of positive feedbacks to warming (due to higher enzyme kinetic rates) could be reversed, leading to lower enzyme production and less feedback to warming: the case of temperature sensitive mortality, and the case of temperature-dependent MGE in an initially warm ecosystem (Supplementary Fig. 5c, d). In all other cases, evolutionary adaptation to warming leads to larger resource allocation to exoenzyme production and a stronger positive feedback, with a greater response in initially colder ecosystems.

### Comparing the ECOS and the EVO responses

The evolutionary response of SOC equilibrium to warming combines the non-evolutionary response of the ecosystem and the evolutionary adaptation of enzyme production (Fig. 1c). In all three scenarios of temperature dependence, the SOC non-evolutionary equilibrium generally decreases as temperature or resource allocation to exoenzymes increases (Fig. 1b, Supplementary Figure 4, Supplementary Note 5). The negative effect of temperature results from the SOC equilibrium being mostly sensitive to the maximum decomposition rate 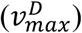 (Supplementary Note 2); all other temperature-dependent parameters have a lesser influence on the SOC equilibrium across their range of variation with temperature. As the maximum decomposition rate increases with warming, the SOC equilibrium decreases. The SOC equilibrium is always lower in systems where microbes invest more resources in exoenzymes, because all other parameters being fixed, a larger enzyme allocation fraction entails that more exoenzymes are produced per unit time, which leads to more SOC decomposed per unit time (Supplementary Note 1). We therefore predict that, without evolution, the equilibrium soil carbon stock shrinks as the climate warms, in all scenarios of temperature-dependence (Fig. 3). The strongest losses occur in initially cold systems due to the non-linear response of the maximum decomposition rate 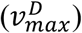 and therefore of SOC equilibrium to temperature (Fig. 3, Supplementary Figure 4). We therefore expect evolution to alter the non-evolutionary response according to the scenario, from aggravating soil carbon loss in systems where microbes adapt to warming with higher exoenzyme production, to buffering carbon loss from soils where microbes adaptively respond to warming with a lower exoenzyme allocation fraction (Fig. 1c).

As expected, the simulated evolutionary response of SOC equilibrium to warming (Figs. 2, 3) mirrors the adaptive response of the enzyme allocation fraction to warming in all scenarios of temperature dependence (Supplementary Figure 5a-d). In the baseline scenario, we showed that microbes always evolve a higher enzyme allocation fraction in response to rising temperature, and we predict that evolution should amplify the non-evolutionary soil carbon loss due to warming (Fig. 3a). The effect of evolution is strongest in ecosystems characterized by conditions that are hostile to microbial growth (low initial temperature, *T*_0_; high mortality, *d_M_*; low MGE, *γ_M_*; low maximum uptake rate, 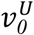) (Fig. 2, Fig. 3a, Supplementary Figure 6), where the adaptive response of the enzyme allocation fraction (change in *φ*\*, equation (2), Supplementary Figure 5a) has been shown to be the most pronounced. Strong evolutionary effects are robust to the other model parameters – enzyme parameters (efficiency, production) and environmental parameters (litter input, leaching) (Supplementary Figures 6 and 7, Supplementary Note 4), which have no influence on the relationship between *φ*\* and temperature (equations (1) and (2)). This relationship, however, cannot explain why stronger evolutionary effects are predicted when the differential accessibility to resources between microbial strains is small (low *c*_0_) (equation (2), Fig. 2c-d). In this case, low competition asymmetry selects for microbes allocating little to exoenzymes (low *φ*\*), shaping systems that are more sensitive to change in enzyme allocation fraction due to the non-linear response of SOC equilibrium to *φ* (Supplementary Note 1, Supplementary Figure 3).

**Figure 2.**
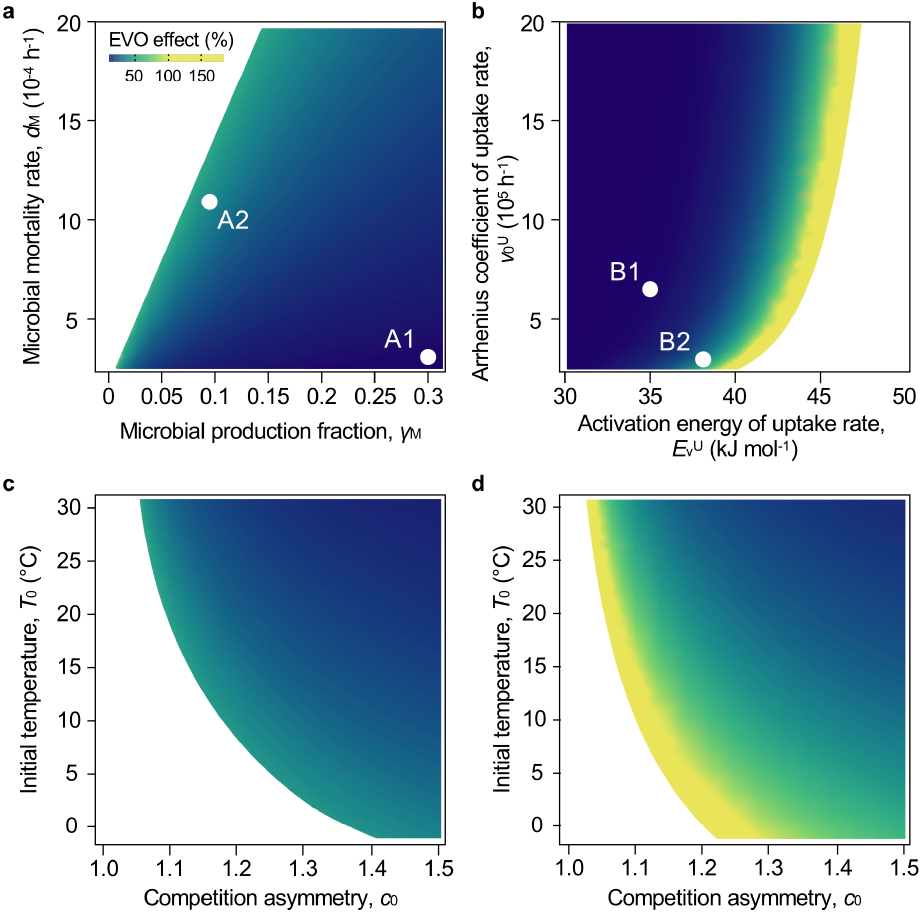
Effect of microbial evolutionary adaptation on the SOC equilibrium response to + 5 °C warming (EVO effect). Temperature influences enzyme kinetics only (baseline scenario of temperature dependence). **a**, Influence of microbial biomass production efficiency, *γ_M_*, and microbial mortality rate, *d_M_*. **b**, Influence of microbial resource acquisition traits 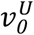 and 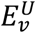. **c-d**, Influence of competition asymmetry, *c*_0_, and initial temperature, *T*_0_. In all figures, constant parameters are set to their default values (Supplementary Table 1) and *I* is set to 5 10^−3^. Points A1 and B1 indicate the default parameter values. Point A2 (respectively B2) exemplifies values of *γ_M_* and *d_M_* (resp. 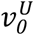 and 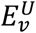) for which the EVO effect is strong. Panel **c** (resp. **d**) shows the influence of *c*_0_ and *T*_0_ on the EVO effect at A2 (resp. B2).

In the temperature-dependent mortality scenario, the strength of the effect of temperature on mortality is an important determinant of the adaptive response to warming. When mortality is moderately sensitive to temperature, the positive response of *φ*\* to warming is attenuated. As a result, the evolutionary aggravation of soil carbon loss is less severe (Fig. 3b). When mortality is strongly sensitive to temperature, warming creates more hostile conditions for microbial growth, therefore *φ*\* decreases with warming. Evolutionary adaptation then buffers the loss of soil carbon (negative EVO effect, Fig. 3c).

**Figure 3.**
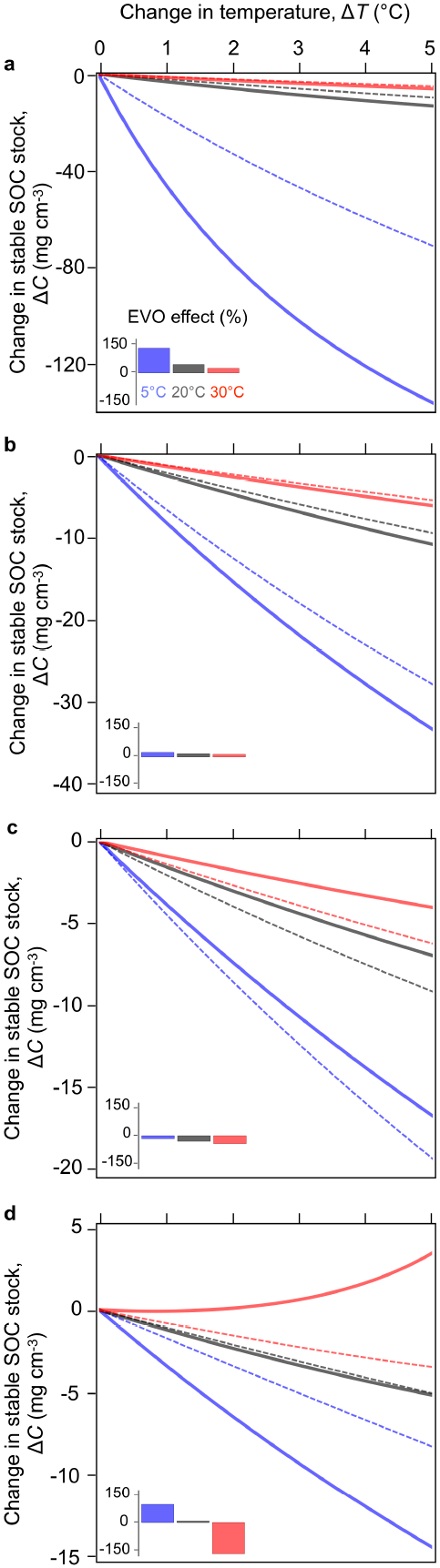
Ecosystem and ecosystem-evolutionary responses of SOC equilibrium to warming (up to + 5 °C) for three scenarios of temperature dependence. Ecosystem and ecosystem-evolutionary changes in SOC equilibrium *C* given by Eq. (3a) (without evolution, dashed curves) and Eq. (3b) (with evolution, plain curves) are plotted as a function of the increase in temperature. *Blue curves*, initial temperature *T*_0_ = 5°C. *Black curves, T*_0_ = *T*_ref_ = 20°C. *Red curves, T*_0_ = 30°C. *Insets*, Direction and magnitude of EVO effect (%), from – 150 % to + 150 %, color code indicates *T*_0_ as before. **a**, Baseline scenario of temperature dependence (enzyme kinetics only). **b**, Temperature-dependent microbial turnover, with 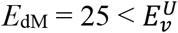. **c**, Temperature-dependent microbial turnover, with 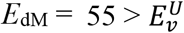. **d**, Temperature-dependent MGE, with *m* = 0.014. Parameters values correspond to point B2 in Fig. 2 (I = 5 10^−3^, 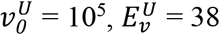, *c*_0_ = 1.17); other parameters are set to their default values (Table S1).

In the temperature-dependent MGE scenario, the direction (positive or negative) of the response of *φ*\* to warming is determined by the initial temperature *T*_0_. The evolutionary effect parallels the response of *φ*\*. At low *T*_0_, *φ*\* increases strongly with temperature, causing an aggravation of soil carbon loss (positive evolutionary effect). In contrast, at high *T*_0_, *φ*\* decreases strongly with warming, thus opposing the non-evolutionary response (negative evolutionary effect) and promoting carbon sequestration instead (Fig. 3d). At intermediate *T*_0_, *φ*\* is barely sensitive to temperature, and the effect of evolutionary adaptation to warming is negligible.

Our general analysis of the ecosystem *CDMZ* model has shown that non-evolutionary response of equilibrium SOC to warming always involves soil carbon loss and may only vary in amplitude. In contrast, the adaptive response of microbial enzyme allocation, which is shaped by an interaction among the non-linear temperature dependences of microbial traits, can drive negative as well as positive responses of equilibrium soil carbon to warming. Adaptive evolution generally amplifies soil carbon loss. However, in ecosystems where soil microbial mortality increases with temperature, this general evolutionary effect can be reversed; adaptive evolution may then reduce soil carbon loss, or even cause carbon sequestration.

### Model predictions using empirical data from five sites

To illustrate how evolutionary effects may vary in real ecosystems, we used available data^22^ on the decomposition kinetic parameters from five sites of increasing latitude and decreasing mean annual temperature (Costa Rica, California, West Virginia, Maine, and Alaska, Fig. 4). We evaluated non-evolutionary and evolutionary responses for each site under three levels of competition asymmetry (as quantified by the local competitive advantage to producers, *c*_0_) (Fig. 4a-h). Under our baseline scenario, EVO effects correlate strongly with mean annual temperature, even more so for low competition asymmetry (Fig. 4i). Stronger EVO effects occur in colder sites, as found in the general analysis (Fig. 3a). In contrast, the non-evolutionary response does not correlate with mean annual temperature (Fig. 4a). As a result, a temperate site such as Maine exhibits a weak non-evolutionary response that can be strongly amplified by evolution, whereas the warm Costa Rica site shows a strong non-evolutionary response that is little affected by evolution.

**Figure 4.**
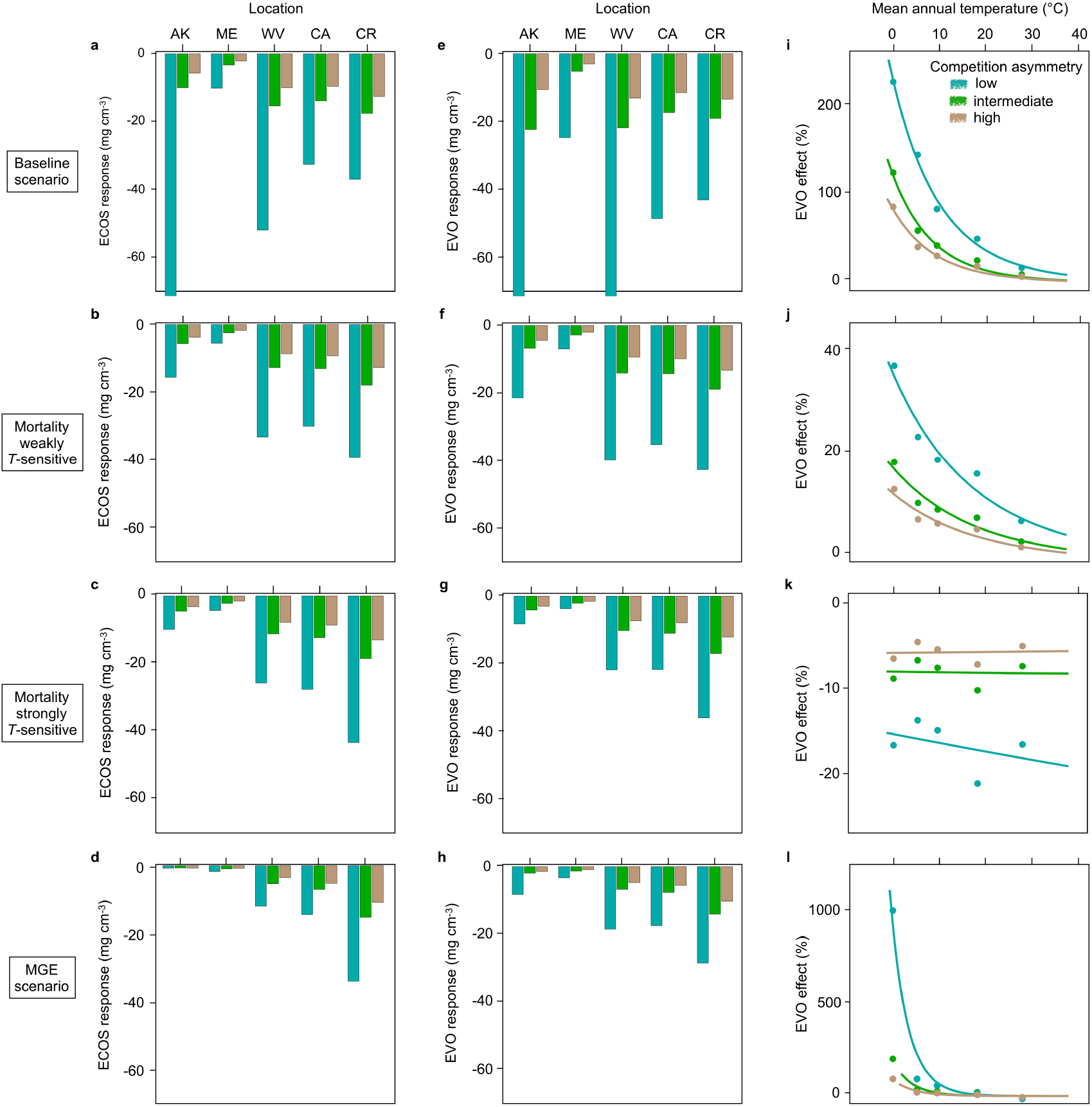
Ecosystem and ecosystem-evolutionary responses of SOC equilibrium to + 5 °C warming predicted for five sites. **a-d**, ECOS response. **e-h**, EVO response. **i-l**, EVO effect. AK: Alaska, boreal forest, *T*_0_ = 0.1°C. ME: Maine, temperate forest, *T*_0_ = 5°C. WV: West Virginia, temperate forest, *T*_0_ = 9°C. CA: California, temperate grassland, *T*_0_ = 17°C. CR: Costa Rica, tropical rain forest, *T*_0_ = 26°C. First row **(a, e, i)**: baseline scenario of temperature dependence. Second row **(b, f, j)**: temperature-dependent microbial turnover scenario with 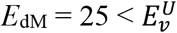. Third row (*c, g, k*): temperature-dependent microbial turnover scenario with 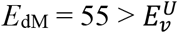. Fourth row (**d, h, l**): temperature-dependent MGE scenario (*m* = – 0.014). The influence of competition asymmetry, *c*_0_, is shown (low: *c*_0_ = 1.17, intermediate: *c*_0_ = 1.34, high: *c*_0_ = 1.5). For clarity, vertical axis for ECOS and EVO responses are truncated at – 65 mg C cm^−3^. Actual values for AK with *c*_0_ = 1.17 are ECOS response = – 170 mg C cm^−3^ and EVO response = – 556 mg C cm^−3^; actual value for WV with *c*_0_ = 1.17 is EVO response = – 92.8 mg C cm^−3^. Parameter values correspond to point B2 in Fig. 2 (*I* = 5 10^−3^, 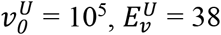, *c*_0_ = 1.17); other parameters are set to their default values (Supplementary Table 1).

These results are quantitatively attenuated but qualitatively unaffected when microbial mortality increases moderately with temperature (Fig. 4b, f, j). With a stronger effect of temperature on microbial mortality, all sites show the evolutionary buffering effect (Fig. 4k) found in the general analysis (Fig. 3c). The intensity of evolutionary buffering is independent of the sites’ mean annual temperature, whereas it varies significantly with competition asymmetry (Fig. 4k). Under the temperature-dependent MGE scenario, non-evolutionary and evolutionary responses are reduced in magnitude compared to the baseline scenario (Fig. 4d, h), particularly in cold sites. However, in these sites, EVO effects are enhanced dramatically (Fig. 4l). Thus, in a site as cold as Alaska, a significant evolutionary loss of soil carbon is predicted, whereas the non-evolutionary-driven loss of soil carbon would be negligible (Fig. 4l).

## Discussion

As global warming increases environmental temperatures, our model predicts evolution of the enzyme allocation fraction, with potentially large effects on the decomposition process and SOC stock (Fig. 1). The size of evolutionary effects is most sensitive to MGE, microbial mortality, activation energy of uptake maximal rate, competition asymmetry, initial temperature (Fig. 2), and to the traits’ temperature sensitivity (Fig. 3). Implications of our findings for large geographic scales across terrestrial ecosystems, are revealed by specifying the model for the five contrasting sites for which exoenzyme kinetics data are available^22^. We find evolutionary aggravation of soil carbon loss to be the most likely outcome, with a strong latitudinal pattern, from small evolutionary effects at low latitude to large evolutionary effects at high latitudes (Fig. 4). Strong temperature-dependence of microbial mortality, however, would dramatically change the evolutionary pattern, possibly causing an attenuation of soil carbon loss or even carbon sequestration in response to warming.

These modeling results are broadly consistent with empirical work showing that changes in soil C stocks are driven by changes in microbial enzyme activity ^35,36^, and that these changes in activity arise from the multiple effects of temperature on enzyme kinetics, microbial pool size, and microbial allocation to exoenzymes^21,37^. Measured mass-specific potential enzyme activity, used as a proxy for allocation to exoenzymes, was shown to generally increase with warming, a pattern also predicted by our evolutionary model^21^. In particular, Steinweg *et al.*^21^ found that the rise in enzyme allocation was greatest for moderate warming and less pronounced for strong warming, which matches our model predictions under the scenario of temperature-dependent MGE. A large body of empirical observations and controlled experiments at various time and spatial scales highlights a general response of soil respiration and soil C stocks to warming, involving an ephemeral increase of respiration, no significant change in SOC, and a decrease in microbial biomass. Only the temperature-dependent mortality scenario with evolution could match these patterns. It has been argued, however, that most experiments remain insufficient to rule out model predictions that depart from these patterns^38^.

The mechanism (physiological, ecological and/or genetic changes) by which enzyme allocation varies in these experiments is unknown; future research should address the role that evolutionary processes may have played in shaping these responses. It is important, in both experimental and modeling studies, to distinguish among different potential biological mechanisms. Next, we consider the distinct roles of physiological acclimation (phenotypic plasticity within individuals) and ecological community assembly (shifts in species composition in whole communities) versus evolutionary adaptation.

Acclimation responses, whereby individuals’ physiological mechanisms buffer the kinetics effect of warming and maintain homeostasis in key life-history traits such as growth and maintenance, are sometimes referred to as ‘adaptation’ (e.g. of microbial carbon use efficiency in response to warming)^18,23,24^. However, this is phenotypic plasticity at the individual level, rather than genetic adaptation at the population level. In our model, constant mortality (in the baseline and temperature-dependent MGE scenarios) and constant MGE (in the baseline and temperature-dependent mortality scenarios) can be interpreted as expressions of such phenotypic plasticity^18,32^. Our results thus show that plasticity, whereby individual cells buffer growth efficiency and/or mortality against temperature variation, does not necessarily impede or overwhelm evolutionary adaptation to climate warming. It is precisely under the assumptions that MGE and mortality remain constant with respect to temperature that the strongest adaptive change in exoenzyme allocation is predicted.

The enzyme allocation fraction itself might be plastic. Indeed, Steinweg *et al.*^21^ interpreted the positive effect of temperature on allocation to enzyme production as a cell-level response to larger nutrient needs driven by higher maintenance costs. In our model, warming causes larger rates of nutrient uptakes (higher 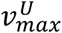) and this can lead to an adaptive increase of the enzyme allocation fraction (equation (1)). From the general evolutionary theory of phenotypic plasticity^39^, we expect the evolutionary optimal reaction norm of enzyme allocation fraction to follow the same pattern, i.e. *φ* increasing with warming, provided that the cost of plasticity is not too high and does not alter too markedly the selection gradient on *φ*.

An interesting way of testing the role of evolutionary adaptation *vs.* plasticity would be to monitor the effect of warming on experimentally evolving bacterial communities at different levels of medium diffusivity or porosity^40^. Our model predicts that the diffusion of resources (DOC), that is likely dependent on physical quality of the soil medium, has a strong influence on the adaptive evolution of microbial exoenzyme production in response to warming, but not on the ecosystem (non-evolutionary) response driven by enzyme kinetics and physiological plasticity. Thus, comparing population-scale decomposition and respiration across treatments that factorially cross temperature and medium quality may contribute to disentangle the effects of adaptation *vs*. plasticity.

An even greater challenge is to tease apart evolutionary adaptation and ecological responses such as species sorting or community shifts in species or functional group abundances, and assess their relative importance in the response of microbial communities to environmental change^25–27^. On the theoretical side, evolutionary and non-evolutionary trait-based models make fundamentally different predictions^41^. Evolutionary models address the dynamics of trait distributions driven in trait space by natural selection and genetic variation. In contrast, ecological community models focus on the dynamics of a given set of species (with possible additions of potentially very different ones via dispersal^42^). In ecological community models, warming may result in a shift of community composition, but only within the set of initially assembled species. As a consequence, different (initial) assemblages may lead to different responses^43^, and the stochasticity of community assembly may therefore contribute to variation in functional responses^44^. In contrast, adaptive evolution fueled by genetic variation (due to *de novo* mutations and/or mobile genetic elements^45^) drives a gradual exploration of the community trait space and convergence towards evolutionarily attractors (‘fitness peaks’). This is expected to yield more predictable community responses, because evolving communities are less constrained by the (historically contingent) set of species and corresponding traits that initially assembled.

On the empirical side, there is growing experimental evidence for soil bacterial communities shifting, with functional consequences on decomposition, in response to climate^43^. On the other hand, evolution experiments on *Pseudomonas* bacteria showed that local adaptation was as important as community composition in shaping the community response to elevated temperature over the course of a two-month experiment^46^. Our model could be extended to disentangle the role of evolutionary adaptation vs. ecological shifts in response to warming. To represent the functional diversity of a microbial community exploiting a diversity of substrates, different SOC pools could be included^47^, alongside specifying different types of enzymes and different microbial functional groups that produce them^20^. In our model parameterization, microbial functional groups would potentially differ in traits such as MGE (*γ_M_*), enzyme cost (*γ_Z_*) and uptake rate 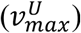^20^, and assuming heritable variation, the evolution of these traits would drive the adaptive response of the microbial functional community to environmental change. In an evolution experiment using *Neurospora discreta* as a model system to assess adaptation of soil fungi to warming^48^, Allison *et al*.^49^ did find evidence for evolutionary responses in MGE and uptake rates. By letting multiple traits evolve, our model extended to multiple substrates could be used first to generate an evolutionarily stable community at a given temperature (following Sauterey *et al.*’s^50^ approach for including evolutionary dynamics in models of ocean planktonic communities), and then evaluate both ecological and evolutionary responses to warming. Ideally, mechanistic models of the kind studied here, without and with genetic variation in exoenzyme allocation and other adaptive traits, could be fitted to experimental data and compared in an inferential model selection framework^51^. Microbial systems offer much promise to achieve such a level of model-data integration. The trait-based approach for microbial communities could facilitate model validation by leveraging genomic and metagenomic data mapped to soil microbial function^52^.

In sum, this work shows that evolutionary feedbacks, operating alongside physiological and ecological responses, may profoundly alter ecosystem function, revealing a critical need for evidence of the role of evolutionary change in microbial communities facing environmental change. Rather surprisingly, studies that investigate how rapid evolution in communities affect ecosystem function are dominated by studies of large organisms, e.g. fish^53^. Rapid evolutionary adaptation of microbial populations to environmental change, especially temperature gradients^54^, is well documented in laboratory systems, but there is limited evidence for the role of positive selection and rapid evolution in the response of natural microbial communities to environmental change^25^. In the laboratory, evolution experiments on single-strain bacterial cultures exposed to a temperature gradient could be analysed by sequencing genes involved in the synthesis of carbon-mineralization enzymes and reconstructing time series of allelic frequency spectra^55^. In such experiments, concurrent measurements of respiration, enzyme activity and microbial biomass could be correlated to putative changes in allele frequencies. A few controlled evolution studies in soil^56,57^, including the resident community, exist for bacteria and viruses, showing patterns of reciprocal (antagonistic) evolution of resistance and infectivity over a matter of weeks. For more complex microbial communities, studies of the human gut are paving the way, suggesting significant short-term evolution occurring in natural microbial populations that are embedded in larger microbial communities^58^. Thus, there is mounting evidence that the diversity and structural complexity of microbial communities does not offset their evolutionary potential to respond to environmental change.

Given the large population size and short generation time of many microorganisms, microbial evolution is likely an essential component of ecosystem response to warming. Microbial evolutionary adaptation to warming, and its impact on the decomposition of soil organic matter, can radically change soil carbon dynamics. Overall, we expect evolutionary effects to vary greatly among ecosystems that differ in biotic (microbial life history and physiology) and abiotic (temperature, soil texture and moisture) characteristics. Empirical data suggest that natural values of enzyme allocation fraction are low^30,34^ and fall in the range for which our model predicts large evolutionary responses of decomposition to warming. Predicting geographic variation in evolutionary responses and effects across large geographic scales calls for more empirical data on variation in ecosystem (competition) traits, especially how the competitive advantage to exoenzyme producers varies with soil physical properties, and in microbial physiological traits, especially microbial mortality. Both the general analysis and the numerical application to specific sites underscore the critical effect of the shapes of the traits’ temperature dependencies. Our work highlights the need empirically to probe the thermal dependence of enzymatic and microbial physiological traits, especially mortality, across wide enough temperature ranges.

In spite of an increasing effort to document and understand the ecosystem impact of microbial physiological and ecological responses to climate warming^18,24,59^, no Earth system model that seeks to represent the role of living organisms in climate feedbacks has yet included evolutionary mechanisms of adaptation. Our model is a critical first step. Next steps will involve accounting for soil carbon stabilization on a timescale longer than respiration^60^; for vegetation types, the associated diversity of organic substrates in litter, and corresponding diversity of microbial decomposers^20^; and for the interaction of biogeochemical cycles and associated stoichiometric constraints^19,20,34,61,62^. As demonstrated by successful Earth-scale modeling of phytoplankton abundance and distribution in the global ocean^63^, future models that take these new steps will trade off their added structural complexity with the ‘self-parameterization’ that the processes of trait heritable variation and natural selection drive in the models themselves. As this research program unfolds, we expect projections of future climate and carbon cycle feedbacks, and their uncertainty, to be significantly impacted by adaptive evolution, from local to global scales.

## Methods

### Ecosystem model

Based on ref. 18^18^ (Fig. 1a, Supplementary Fig. 1), the ecosystem model has four state variables measured in unit mass of carbon: soil (non decomposed) organic carbon (SOC), *C*; soil decomposed soluble organic carbon (DOC), *D*; microbial biomass, M; and exoenzyme concentration, *Z*. Exoenzyme production drives the decomposition process of *C* into *D*, which is the only source of carbon for microbes. The model accounts for microbial production and death, exoenzyme decay, recycling of dead microbes and degraded exoenzymes, *C* input from plant litter, and leaching of *C* and *D*.

#### Model equations

State variables *C, D, M, Z* obey equations (1a–d):

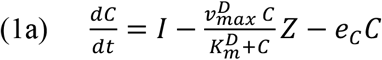

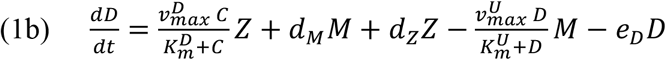

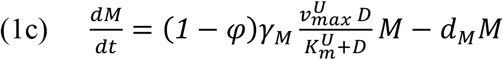

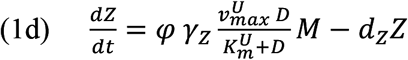

In equation (1a), decomposition follows from Michaelis-Menten kinetics of *Z* binding substrate *C*; there is a constant input, *I*, of soil organic (non decomposed) carbon from aboveground litter, and a loss due to leaching at constant rate *e_C_*. In equation (1b), *D* is produced by decomposition and the recycling of dead microbial biomass and inactive enzymes; *D* is consumed by microbial uptake, and lost by leaching at constant rate *e_D_*. In equation (1c), growth of microbial biomass *M* is driven by the rate of *D* uptake (a Monod function of *D*) times the fraction of uptaken *D* turned into biomass, (1 – *φ*) *γ_M_*, minus microbial mortality at constant rate *d_M_*. In equation (1d), enzyme variation is driven by the rate of *D* uptake times the fraction allocated to enzyme production, *φ*, and production efficiency, *γ_Z_*, minus enzyme deactivation at constant rate, *d_Z_*.

#### Ecosystem equilibria

The ecosystem model possesses either one globally stable equilibrium, or three equilibria (one of which is always unstable) (Supplementary Fig. 2). There are thresholds *φ*_min_ and *φ*_max_ such that the single globally stable equilibrium exists for *φ* < *φ*_min_ or *φ* > *φ*_max_ and is given by *C* = *I*/*e_C_*, *D* = 0, *M* = 0, *Z* = 0. Thus, at this equilibrium, the microbial population is extinct and no decomposition occurs. For *φ*_min_ < *φ* < *φ*_max_, the microbial population can either go extinct (then the system stabilizes at the same equilibrium as before) or persists at or around a non-trivial equilibrium, which can be solved for analytically (equation (S1)). Note that φmin and φmax depend on all microbial and model parameters (Supplementary Figure 3, Supplementary Note 1).

#### Effect of temperature on model parameters

Decomposition is predicted to respond to warming^5^ due to the temperature sensitivity of enzymatic activity^22,29,64^. Microbial assimilation may also vary with temperature if the microbial membrane proteins involved in nutrient uptake are sensitive to warming. Following ref. 18^18^, we assume that exoenzyme kinetics parameters (maximum decomposition rate 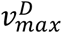 and half-saturation constant 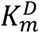) and microbial uptake parameters (maximum uptake rate 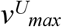 and half-saturation constant 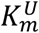) follow Arrhenius relations with temperature. This defines our baseline ‘kinetics-only’ scenario of temperature-dependent decomposition:

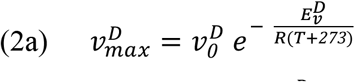

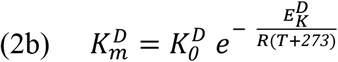

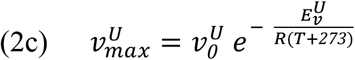

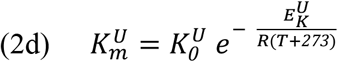

where *T* is temperature in Celsius, *R* is the ideal gas constant, and the *E* parameters denote the corresponding activation energies.

We consider two additional scenarios for the influence of temperature on decomposition. In the temperature-dependent microbial mortality scenario^32^, the microbial death rate increases with temperature. This could be due to a higher risk of predation or pathogenic infection at higher temperatures, or faster microbial senescence due to higher protein turnover^32^. In this scenario, the microbial death rate dM depends on temperature according to

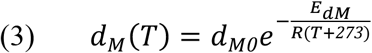

as in ref. 32^32^.

In the temperature-dependent microbial growth efficiency (MGE) scenario, the MGE decreases with temperature^18,24,32^, possibly due to higher maintenance costs at higher temperature^33^. This is modeled by making the microbial growth efficiency γM vary linearly with temperature^18,22,47,65^:

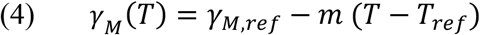

with *T*_ref_ = 20 °C.

How scenarios of temperature-dependence and parameter values influence the response of equilibrium *C* to temperature is shown in Supplementary Figure 4 and commented on in the Supplementary Note 4.

### Evolutionary Dynamics

The enzyme allocation fraction *φ* is a ‘public good’ trait: as an individual microbe produces exoenzymes, it experiences an energetic cost and obtains a benefit – access to decomposed organic carbon – that depends on its own and other microbes’s production in the spatial neighborhood^66,67^. As a public good trait, *φ* is under strong direct negative selection: ‘cheaters’ that produce less or no exoenzymes, and thus avoid the cost while reaping the benefit of enzyme production by cooperative neighbors, should be at a selective advantage. In a highly diffusive environment in which exoenzymes are well mixed, *φ* would evolve to zero, leading to evolutionary suicide^68^. However, in a more realistic spatially distributed environment with limited exoenzyme diffusion, microbes with a given trait are more likely to interact with phenotypically similar microbes, which puts more cooperative microbes at a competitive advantage over less cooperative strains^66,69,70^. This generates indirect positive selection on trait *φ*.

We derived the trait value *φ*\* at which negative and positive selections balance, giving the evolutionarily stable microbial strategy (Supplementary Note 3), as

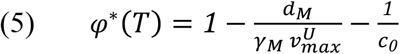

where *c*_0_ measures the competitive advantage, due to spatially local interactions, of any given strain over a slightly less cooperative strain, or ‘competition asymmetry’ (Supplementary Note 3). The parameter *c*_0_ is likely to depend on the diffusivity of *D*, which may itself vary with soil properties such as texture or water content.

As temperature rises from *T*_0_ to *T*, the direction and magnitude of the microbial adaptive response is measured by Δ*φ*\* = *φ*\*(*T*) – *φ*\*(*T*_0_), which depends on the scenario of temperature dependence. The evolutionary response (EVO response) of SOC is given by

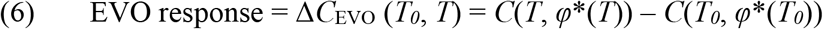

where *C*(*T, φ*) denotes ecosystem equilibrium *C* at temperature T, given enzyme allocation fraction *φ*. The EVO response is to be compared with the response in the absence of evolution (ECOS response):

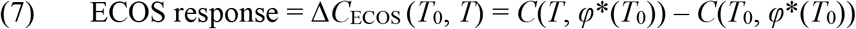

in which the enzyme allocation fraction is fixed at its *T*_0_-adapted value, *φ*\*(*T*_0_) (Fig. 1c).

We measure the magnitude of the evolutionary effect (EVO effect) as the difference between the EVO response averaged over the temperature range (*T*_0_, *T*) and the ECOS response averaged over the same temperature range, normalized by the ECOS response:

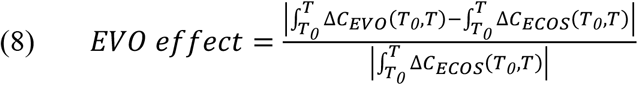

This evaluation allows us to compare EVO effects across systems that differ in the magnitude of their ECOS response. In all simulations we use *T* = *T*_0_ + Δ*T* where Δ*T* = 5 °C. In general, the ECOS and EVO responses are monotonic, close-to-linear functions of *T* over the considered temperature ranges (*T*_0_, *T*_0_ + Δ*T*), which makes all our comparative analyses almost insensitive to our choice of Δ*T*.

### Parameter default values and sensitivity analysis

Under the temperature-dependent kinetics-only scenario, the ecosystem model (equations (1) and (2)) includes seven microbial parameters 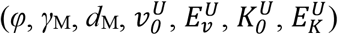, six enzyme parameters 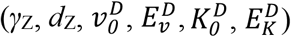 and four environmental parameters (*I, e_C_, e_D_, T*). Our set of default parameter values is derived from Allison *et al.* (2010)^18^ (Supplementary Table 1). The enzyme allocation fraction default value is 10% at 20 °C^34^. For the dependence of enzyme kinetics parameters on temperature, 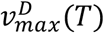 and 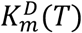, we selected the Arrhenius equations that best fit data from California^22^ (mean annual *T* = 17 °C) and match values at 20 °C (0.42 and 600, respectively)^18^. For the uptake kinetic parameters, we obtained 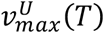 by selecting 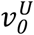 that best fits the Arrhenius equation in ref. 18^18^ with 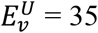 and we obtained 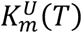 by selecting 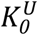 and 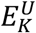 that best fit the linear relation used in ref. 18^18^. To parametrize the temperature-dependent mortality (*d*_M,ref_, *T*_ref_) and MGE (*γ*_M,ref_, *m*, *T*_ref_) models, we used values from ref. 32^32^ and tested two values of *E*_dM_ (*E*_dM_ = 0 is the enzyme only temperature-dependent model). For greater realism, we used a higher value of the exoenzyme deactivation rate (twice the value used in ref. 18^18^) and constrained the range of all parameters in order to enhance stability and produce relative stock sizes that are consistent with empirical data, so that at equilibrium *M* is about 1% of *C*, *Z* is about 1% of *M* and *D* is limiting (hence close to 0 at equilibrium^71,72^),

We analysed the model sensitivity by varying parameters over two orders of magnitude (as in ref. 18^18^) – except *γ_M_* and *γ_Z_* for which we used the whole range over which the nontrivial ecosystem equilibrium is stable (Supplementary Table 2). To assess the significance of our findings for real ecosystems, we focused on five sites for which empirical data^22^ could be used to constrain the model. The five sites contrast strongly in their initial temperature, *T*_0_, and decomposition kinetics, 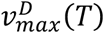 and 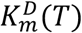 for which we selected the Arrhenius equations (2) that best fit the relations used in ref. 22^22^ (Supplementary Table 3).

## Supporting information

Supplementary Information version 2

## Data availability

No datasets were generated or analysed during the current study.

## Code availability

The sensitivity analysis and figures that support the findings of this study were coded on Mathematica. The code file has been deposited in “MicFigCode”, https://github.com/elsaabs/MicFigCode/tree/master.

## End Notes

## Acknowledgements

We thank Rachel Gallery, Pierre-Henri Gouyon, Moira Hough, Hélène Leman, Laura Meredith, and Mitch Pavao-Zuckerman for discussion. E.A. was supported by fellowships from Ecole Doctorale Frontières du Vivant and MemoLife Laboratory of Excellence (PIA-10-LBX-54). S.R.S. was supported by a grant from the Genomic Science Program of the United States Department of Energy (DOE) (DE-SC0016440) and the University of Arizona’s Agnese Nelms Haury Program in Environment and Social Justice. R.F. acknowledges support from FACE Partner University Fund, CNRS Mission pour l’Interdisciplinarité, and PSL University (IRIS OCAV and PSL-University of Arizona Mobility Program).

## Author contributions

R.F. conceived the study. All authors developed the model. E.A. performed the analysis. E.A. and R.F. wrote the first version of the manuscript. All authors finalized the paper.

## Competing interests

The authors declare no competing interests.

## Additional information

**Supplementary information** is available for this paper at https://doi.org/xxx

**Correspondence and requests for materials** should be addressed to E.A. or R.F.

