## Supplementary Information version 2 for "Microbial evolution reshapes soil carbon feedbacks to climate change"

**This PDF file includes:**

Supplementary Notes
Supplementary Figures 1 to 10
Supplementary Tables 1 to 3
References for SI reference citations

**<u>Supplementary Note 1</u>: Analysis of ecosystem model at constant temperature**

At any given temperature $T$, depending on the enzyme allocation fraction $\varphi$, the ecosystem model possesses either one globally stable equilibrium, or three equilibria (one of which is always unstable) (Supplementary Fig. 3).

The single (globally stable) equilibrium exists for $\varphi < \varphi_{\min}$ or $\varphi > \varphi_{\max}$ and is given by $M = 0$, $Z = 0$, $C = I/e_C$, $D = 0$. Thus, at this equilibrium, the microbial population is extinct and no decomposition occurs.

For $\varphi_{\min} < \varphi < \varphi_{\max}$, the microbial population can either go extinct (then the system stabilizes at the same equilibrium as before) or persists at or around a non-trivial equilibrium, which can be solved for analytically:

(S1a) $C = \frac{1}{e_C} \frac{\alpha - \sqrt{\alpha^2 - 4\beta}}{2\mu}$

(S1b) $D = \frac{d_M K_m^U}{\lambda}$

(S1c) $M = \frac{\gamma_M (1-\phi)}{d_M \delta} \frac{\left(\alpha - \Delta_M + \sqrt{\alpha^2 - 4\beta}\right)}{2\mu}$

(S1d) $Z = \frac{\gamma_Z \phi}{d_Z \delta} \frac{\left(\alpha + \Delta_Z + \sqrt{\alpha^2 - 4\beta}\right)}{2\mu}$

where

$$\alpha = \mu\left(I - e_C K_m^D\right) + \phi \gamma_Z v_{max}^D \varrho$$
$$\beta = \mu \lambda \delta d_Z I e_C K_m^D$$
$$\mu = \left(\phi \gamma_Z v_{max}^D - d_Z \delta\right) \lambda$$
$$\Delta_M = 2 d_Z \delta \varrho$$
$$\Delta_Z = - 2 d_Z \delta \varrho$$
$$\lambda = (1 - \phi)\, \gamma_M v_{max}^U - d_M$$
$$\delta = 1 - (1 - \phi)\, \gamma_M - \phi \gamma_Z$$
$$\varrho = e_C K_m^D \lambda - e_D K_m^U d_M \ .$$

Although an analytical study of the stability of the non-trivial equilibrium is out of reach, numerically we observed stability for most parameter values, based on the calculation of the

Jacobian eigenvalues. Only for $\varphi$ very close to $\varphi_{\min}$ we detected a few cases of equilibrium instability, in which case the system converges toward a small limit cycle. $\varphi_{\min}$ therefore provides a good approximation for the lower bound of the range of $\varphi$ over which the system persists at a non-trivial equilibrium.

Boundary values, $\varphi_{\min}$ and $\varphi_{\max}$, of existence of the non-trivial equilibrium are found where the unstable non-trivial equilibrium meets the stable non-trivial equilibrium, *i.e.* when $\alpha^2 - 4\beta = 0$, that we solved numerically. Empirical data suggest that natural values of $\varphi$ are small[1,2]. The model shows that as $\varphi$ decreases toward $\varphi_{\min}$, equilibrium $C$ becomes more sensitive to $\varphi$, and the smaller $\varphi_{\min}$, the stronger the sensitivity of equilibrium $C$ to $\varphi$ near $\varphi_{\min}$. This is true across the whole parameter set (Supplementary Fig. 3). Hereafter, unless stated otherwise, we simply use $C$ (respectively $D$, $M$, $Z$) to denote equilibrium values at the non-trivial stable equilibrium (Supplementary Equation (1)).

We performed a simple sensitivity analysis of $C$ and existence boundaries $\varphi_{\min}$ and $\varphi_{\max}$ (Supplementary Table 2), following the scheme used by Allison *et al.* (2010)[3] (same range for each parameter in common between our study and theirs, same measure of sensitivity). All parameters affect $\varphi_{\min}$ and $\varphi_{\max}$, whereas only microbial growth efficiency ($\gamma_M$), enzyme allocation fraction ($\varphi$), and enzyme parameters ($\gamma_Z$, $d_Z$, $v_0^D$, $K_0^D$, $E_v^D$, $E_K^D$) have a significant effect on $C$ (Supplementary Fig. 3). This reflects the fact that SOC decomposition is governed by the size of the microbial pool (mostly sensitive to $\gamma_M$) and enzyme pool (shaped by $\varphi$ and $\gamma_Z$), and by enzyme performance (determined by the other enzyme parameters) (Supplementary Table 2).

The lack of sensitivity of $C$ to $d_M$ is intriguing, because $d_M$ has a relatively strong influence on the microbial pool (Supplementary Table 2). Both Allison *et al.* (2010) and Hagerty *et al.* (2014)[4] found a significant effect of $d_M$ on $C$. This difference arises from the process driving the enzyme production. In Allison *et al.* (2010) and Hagerty *et al.* (2014), the enzyme production rate, $r_Z$ (Supplementary Fig. 1b), is constant over time ('constitutive production'). Using notations similar to ours, their model is

$$\frac{dC}{dt} = I_C - \frac{v_{max}^D C}{K_m^D + C} Z + p d_M M$$

$$\frac{dD}{dt} = I_D + \frac{v_{max}^D C}{K_m^D + C} Z + (1-p) d_M M + d_Z Z - \frac{v_{max}^U D}{K_m^U + D} M$$

$$\frac{dM}{dt} = CUE \frac{v_{max}^U D}{K_m^U + D} M - d_M M$$

$$\frac{dZ}{dt} = r_Z M - d_Z Z$$

where $CUE$ is carbon use efficiency (*sensu* Allison *et al.* 2010) and $p$ is the fraction of microbial necromass recycled in SOC *vs*. DOC. When solving this system at equilibrium, the enzyme concentration, $Z$, only depends on enzyme traits ($\gamma_M$ and $\gamma_Z$) and microbial abundance, $M$ ( $Z = \frac{r_Z}{d_Z} M$ ), which depends on microbial turnover rate ( $M = \frac{CUE\,(I_D + I_C)}{(d_M + r_Z)(1 - CUE)}$ ). As a consequence, equilibrium $Z$ is negatively dependent on $d_M$, therefore equilibrium $C$ is positively dependent on $d_M$. The analytical expression of equilibrium $C$[5] is:

$$C = \frac{-d_Z K_m^D (I_C (d_M (1+CUE(p-1)) + r_Z (1-CUE)) CUE I_D p d_M)}{I_C (d_M (d_Z (1+CUE(p-1))) + r_Z (d_Z (1-CUE) - CUE v_{max}^D)) + CUE I_D (p d_M d_Z - r_Z v_{max}^D)}$$

To link exoenzyme production per individual microbe to the amount of assimilated resource, our model (equation (1)) substitutes *CUE* with $\varphi CUE$ and $r_Z$ with $\phi CUE \frac{v_{max}^U D}{K_m^U + D}$ . With exoenzyme production being a fraction of assimilated resource, *M* at equilibrium is directly inversely proportional to $d_M$:

$$M = \frac{(1-\phi) CUE (I_D + I_C)}{d_M (1-CUE)}$$

and *Z* becomes independent of $d_M$:

$$Z = \frac{\phi d_M}{d_Z (1-\phi)} M = \frac{\phi CUE (I_D + I_C)}{d_Z (1-CUE)} .$$

As a consequence, *C* does not depend on $d_M$ and is given by:

$$C = \frac{-d_Z K_m^D (I_C (1+CUE(p(1-\phi)-1)) + CUE I_D p (1-\phi))}{I_C (d_Z (1+CUE(p(1-\phi)-1))) + CUE I_D (p d_Z (1-\phi) - v_{max}^D \phi)} .$$

This expression was obtained in the case where the production costs of microbial biomass and exoenzymes are the same ($\gamma_Z = \gamma_M$ = CUE). Including leaching of *D* and *C* re-establishes a dependency of equilibrium *C* on $d_M$, but the sensitivity to $d_M$ remains very low across values of leaching rates for which the equilibrium exists.

To summarize, in models where exoenzymes are produced at a constant rate per unit microbial biomass, the equilibrium exoenzyme concentration, hence the SOC equilibrium stock, are strongly dependent upon all microbial parameters, including $d_M$. In models like ours where exoenzyme production depends on the amount of assimilated resource, the equilibrium exoenzyme concentration and SOC stock only depend on the allocation parameters, enzyme parameters and resource input parameters, and not on $d_M$. However, taking evolution into account may restore a dependency of SOC equilibrium on microbial mortality, because *C* is strongly sensitive to the enzyme allocation fraction, $\varphi$, and the evolutionarily stable trait value $\varphi^*$ itself is strongly sensitive to $d_M$ (equation (5) in Methods).

**Supplementary Note 2: Effect of temperature on SOC equilibrium, *C***

For a given enzyme allocation fraction $\varphi$, $C$ always decreases as temperature increases in the baseline scenario (Fig. 1b, Supplementary Fig. 4). In this scenario, the sensitivity of $C$ to temperature is mediated chiefly by the temperature-dependence of enzyme kinetics parameters $v_{max}^{D}$ and $K_{m}^{D}$ (Supplementary Table 2). As $T$ increases, both $v_{max}^{D}$ and $K_{m}^{D}$ increase, causing antagonistic effects on $C$; however, based on empirical data[6], most of the effect of warming is on $v_{max}^{D}$, which makes the thermal effect mediated by $v_{max}^{D}$ prevail. As a consequence, the decomposition rate rises with temperature, hence $C$ declines.

In the temperature-dependent microbial turnover scenario, the effect of temperature on decomposition and SOC stock is virtually unchanged compared to the baseline scenario. This is because the sensitivity of decomposition rate and $C$ to microbial mortality $d_{\mathrm{M}}$ is very small (Supplementary Table 2).

The loss of soil C with warming also holds in the temperature-dependent MGE scenario. In this case, increasing $T$ has antagonistic effects on decomposition mediated by $v_{max}^{D}$ (enzyme activity increases with temperature) *vs.* $\gamma_{\mathrm{M}}$ (microbial growth decreases with temperature). However, the effect of the exponential dependence of $v_{max}^{D}$ on $T$ is stronger than the effect of the linear dependence of $\gamma_{\mathrm{M}}$ on $T$; the former thus dominates the effect of $T$ on $C$, but the loss of soil C is attenuated compared to the baseline and the temperature-dependent microbial turnover scenario.

In all three scenarios, $C$ is more sensitive to $T$ across lower values of $T$ (Fig. 3, Supplementary Fig. 4) because $C$ is most sensitive to and a function of $-v_{max}^{D}$, which sensitivity to $T$ decreases as $T$ increases. As a consequence, the loss of soil C with warming is more pronounced in colder ecosystems.

### <u>Supplementary Note 3</u>: Evolutionary model

We use the framework of adaptive dynamics[7,8]. In this framework, evolution is modeled as a competition process between a ‘resident strategy’ (wild-type) and alternate strategies (mutants) within a set of feasible phenotypes. In a given environment (*e.g.* at a given temperature), an ‘adaptation’ or ‘adapted value’ of a trait is a phenotype that (i) when resident, no mutant can invade; (ii) can be reached by a sequence of phenotypic substitutions, whereby each step involves the replacement of a resident phenotype by a mutant phenotype. Here the phenotypic trait of interest is the enzyme allocation fraction, $\varphi$. The set of feasible phenotypes is the range ($\varphi_{\mathrm{min}}$ , $\varphi_{\mathrm{max}}$) at a given temperature, for which the non-trivial ecosystem equilibrium exists.

*Interaction between resident and mutant strains*

To model the competition effect of a resident phenotype, $\varphi_{\mathrm{res}}$, on the population growth of a mutant phenotype, $\varphi_{\mathrm{mut}}$, we extend the ecosystem model written for a single type (Equation (1) in Methods). To account for the local nature of the interaction between rare mutant and common resident cells, we introduce a function (hereafter denoted by *c*) of the difference between $\varphi_{\mathrm{res}}$ and $\varphi_{\mathrm{mut}}$ to measure how local decomposition by mutant and resident cells differ from ‘mean field’ (average) decomposition by resident cells. Thus, for given *C*, *D*, *Z*, the growth of the mutant population is governed by

$$\text{(S2)} \qquad \frac{dM_{mut}}{dt} = (1-\phi)\,\gamma_M \frac{v^U_{max}(1+c(\phi_{mut}-\phi_{res}))D_{res}}{K^U_m+(1+c(\phi_{mut}-\phi_{res}))D_{res}} M_{mut} - d_M M_{mut}$$

where $D_{\mathrm{res}}$ is the equilibrium *D* predicted by the ecosystem model applied to the sole resident phenotype $\varphi_{\mathrm{res}}$. Here function *c* satisfies $c(0) = 0$, $c(z) > 0$ if $z > 0$ and $c(z) < 0$ if $z < 0$.

The underlying assumption is that each microbe has access to DOC partly as a public good and partly as a private good[9]. The public good part results from the diffusion of exoenzymes. The private good part results from local decomposition at the microscopic scale of cells and exoenzymes that they produce themselves. A mutant cell that invests more (resp. less) in exoenzyme has access to more (less) DOC than the average resident cell because the cell’s private good is greater (smaller) whereas all cells share the same public good. In a spatially implicit model like ours, diffusion is not directly modeled, but its effect on the accessibility of DOC to a mutant strain can be phenomenologically accounted for by a parameterizaton that puts mutant cells at a competitive advantage for DOC if the mutant phenotype invests more in exoenzyme production than the resident phenotype , or at a competitive disadvantage if the mutant phenotype invests less . This parameterization is achieved with the function *c* in equation (S2), where $c<1$ when $\varphi_{\mathrm{mut}} < \varphi_{\mathrm{res}}$ and $c>1$ when $\varphi_{\mathrm{mut}} > \varphi_{\mathrm{res}}$.

*Invasion fitness and selection gradient*

Mutant fitness $s(\phi_{mut},\, \phi_{res})$ is given by the mutant population growth rate per unit biomass:

$$\text{(S3)} \qquad s(\phi_{mut},\, \phi_{res}) = (1-\phi)\,\gamma_M \frac{v^U_{max}(1+c(\phi_{mut}-\phi_{res}))D_{res}}{K^U_m+(1+c(\phi_{mut}-\phi_{res}))D_{res}} - d_M$$

The selection gradient then obtains by taking the first order derivative of the invasion fitness with respect to the mutant trait:

(S4) $$\nabla s(\phi) = \frac{d_M}{1-\phi}\left(\left(1 - \phi - \frac{d_M}{v_{max}^U \gamma_M}\right) c_0 - 1\right)$$

where $c_0 = c'(0)$ measures the local competitive advantage to stronger exoenzyme producers, which we call 'competition asymmetry'. Note that by definition of function $c$, we always have $c_0 > 0$. Variation in $c_0$ may be caused by different soil diffusion properties, due to *e.g.* physical texture or moisture.

*Evolutionary singularity*

Trait values that nullify the selection gradient are called 'evolutionary singularities'. An evolutionary singularity can be attractive or repelling, and invadable or non-invadable. Evolutionary singularities that are attractive and non-invadable represent potential end-points of evolutionary adaptation. Evolutionary singularities that are attractive and invadable can lead to evolutionary branching[8].

In a given environment (fixed parameters, constant temperature) there is at most one evolutionary singularity given by defining $\phi^*$ as the value of $\phi$ that makes $\nabla s(\phi) = 0$ in eqn S4:

(S5) $$\phi^* = 1 - \frac{d_M}{v_{max}^U \gamma_M} - \frac{1}{c_0}.$$

Existence of $\varphi^* > 0$ requires $\frac{d_M}{v_{max}^U \gamma_M} < 1$ and $c_0 > \frac{1}{\left(1 - \frac{d_M}{v_{max}^U \gamma_M}\right)}$. Thus, the (cooperative) trait $\varphi$ can evolve above zero only if the local competition advantage to stronger enzyme producers is large enough. The condition for $\varphi^*$ to be evolutionarily stable is $c''(0) < 2c_0^2$ and no other condition than existence is required for $\varphi^*$ to be always convergent. Here we assume that function $c$ is such that $\varphi^*$ is evolutionarily stable and attractive.

Supplementary Equation (5) shows that more cooperation (larger $\varphi^*$) should evolve in microbial populations with lower mortality, greater nutrient uptake, and/or higher MGE. When comparing microbial populations with similar life-history traits $\gamma_M$, $v_{max}^U$ and $d_M$, stronger competitive advantage to exoenzyme producers (i.e. higher $c_0$) selects for larger $\varphi^*$.

*Effect of temperature on $\varphi^*$*

Supplementary Equation (5) also predicts the evolutionary effect of temperature variation on enzyme production. The evolutionary adaptive response of enzyme allocation fraction $\varphi$ to warming is driven by the effect of temperature on $v_{max}^U$, $d_M$, and $\gamma_M$. The thermal dependence of $v_{max}^U$ is assumed, in our baseline scenario, as a consequence of temperature-dependent enzyme kinetics. In this scenario, $d_M$ and $\gamma_M$ are constant (temperature independent); the evolutionary effect of $T$ on $\varphi^*$ is thus entirely mediated by the effect of $T$ on $v_{max}^U$. Warming drives $v_{max}^U$ up, which causes $\varphi^*$ to rise (Supplementary Fig. 5a). The sensitivity of $\varphi^*$ to temperature is strongest at low initial temperature $T_0$ (Supplementary Fig. 5a).

In the temperature-dependent turnover scenario, as the influences of $d_M$ and $v_{max}^U$ on $\varphi^*$ are antagonistic (Supplementary equation (5)), we can analytically determine the direction and amplitude of the effect of temperature on $\varphi^*$ using its its first order derivative with respect to $T$:

$$\frac{d\phi^*}{dT} = \frac{d_M(T)}{v_{max}^U(T)\gamma_M R(T+273)^2}\left(E_v^U - E_{dM}\right)$$

Warming causes $\varphi^*$ to rise ($d\varphi^*/dT > 0$) when ${E_v}^U > E_{dM}$, *i.e.* when $d_M$ is not too strongly sensitive to temperature. As a consequence, the rise of $\varphi^*$ is strongest in the baseline scenario (where $E_{dM} = 0$) (Supplementary Fig. 5a, b). Warming causes $\varphi^*$ to decline when ${E_v}^U > E_{dM}$ (Supplementary Fig. 5c). The influence of $T_0$ on the sensitivity of $\varphi^*$ to temperature observed in the baseline scenario is lost in the temperature-dependent turnover scenario (Supplementary Fig. 5c).

In the temperature-dependent MGE scenario, warming drives $v_{max}^U$ up and $\gamma_M$ down, therefore the strongest rise of $\varphi^*$ occurs if $\gamma_M$ remained constant. Due to the interaction between one trait that responds linearly ($\gamma_M$) and one trait that responds non-linearly to temperature ($v_{max}^U$), both the direction and amplitude of the influence of $T$ on $\varphi^*$ strongly relies on the system initial temperature, $T_0$:

$$\frac{d\phi^*}{dT} > 0 \Leftrightarrow -m + \frac{E_v^U}{R(T+273)^2}\gamma_M(T) > 0 \Leftrightarrow T^2 + T\left(\frac{E_v^U}{R} + 2\times 273\right) + \left(273^2 - \frac{E_v^U\gamma_{M,ref}}{mR} - \frac{E_v^U T_{ref}}{R}\right) < 0$$

Warming causes $\varphi^*$ to rise for values of $T$ that are inside the solutions of the above quadratic equation, which are equal to - 5000°C and 23°C with the parameter values used in Figure 3 and Supplementary Figure 5. As a consequence, in response to warming, $\varphi^*$ rises in systems initially below 23°C, $\varphi^*$ decreases in systems initially above 23°C, and $\varphi^*$ is almost not sensitive to temperature in systems initially around 23°C (Supplementary Fig. 5d).

Quantitatively, the strongest warming effect on $\varphi^*$ is observed in the baseline scenario in cold ecosystems.

### <u>Supplementary Note 4</u>: Sensitivity analysis of ECOS and EVO responses and EVO effects

In the baseline scenario of temperature dependence, the EVO effect is most sensitive to microbial life-history traits $\gamma_M$, $d_M$ (Fig. 2a, Supplementary Fig. 6a,b), $v_0^U$ and $E_v^U$ (Fig. 2b, Supplementary Fig. 6c, d) and environmental parameters $c_0$ and $T_0$ (Fig. 2c-d, Supplementary Fig. 6e, f). This is because these parameters influence the sensitivity of the adaptive strategy, $\varphi$*, to temperature and/or the initial adapted value, $\varphi^*(T_0)$ (equation (S5)), and therefore they can affect the EVO response more than the ECOSresponse, with strong EVO effects as a result (Supplementary Fig. 6a-f).

In contrast, the EVO effect is little affected by the other model parameters: enzyme parameters (efficiency, production) and environmental parameters (litter input, leaching) (Supplementary Figs. 6g-o and 7). Considering enzyme parameters, we find that less efficient enzymes lead to larger ECOSand EVO responses to warming (Supplementary Fig. 6g1-l1). However, the ECOSand EVO responses exhibit similar sensitivities to most enzyme parameters, which therefore have very little impact on the EVO effect (Supplementary Figs. 4a-c and 6i-l). Only the activation energies ( $E_v^D$ and $E_K^D$ ) have a slightly stronger impact on the ECOSresponse than on the EVO response, which results in slightly weaker EVO effects in systems with less efficient enzymes (Supplementary Figs. 6g, h and S7a, b). Finally, environmental parameters – litter input $I$, and leaching rates $e_C$ and $e_D$ – primarily affect the stability of the ecosystem equilibrium (via their influence on $\varphi_{min}$, Supplementary Fig. 3) and have little influence on the EVO effect (Supplementary Figs. 6m-o and 7d).

In the other scenarios of temperature dependence, the influence of parameters on the EVO effect is consistent with results shown in Fig. 2 (sensitivity analysis in the baseline scenario) and Fig. 3 (ECOSand EVO responses and EVO effect for the default system in the other scenarios). Thus, EVO effects are generally strong in cold ecosystems harboring communities of slow-growing microbes, under soil conditions that give only a small competitive edge to greater enzyme producer (such as in Fig. 2); EVO effets are often reduced in systems with stronger dependence of mortality $d_M$ on temperature; and EVO effects are strong in warm ecosystems when MGE decreases with warming, as seen in Fig. 3 (Supplementary Fig. 8).

**<u>Supplementary Note 5</u>: Ecosystem responses to warming**

In absence of microbial evolution, the model predicts a large loss of soil carbon with warming in systems harboring low-productivity microbes (low $\gamma_M$) secreting low-performance enzymes (high $\gamma_Z$); and in systems of high-productivity microbes (high $\gamma_M$) that synthesize high-performance enzymes (low $\gamma_Z$) but invest very little resource in their production ($\varphi$ close to the minimum viable value, $\varphi_{min}$) – which is the expected adaptation to highly diffusive soils (according to Supplementary Equation (5)). The prediction of large ecology-driven loss of soil carbon holds even if the microbial turnover increases with warming, but is attenuated (though not reversed) if MGE decreases with warming. These results are consistent with previous ecosystem models of soil microbial decomposition that compared constant versus decreasing MGE with warming[5,10–13]. Note, in these models, a constant MGE (CUE in models ignoring exoenzymes dynamics, as in German *et al.* (2012)) was viewed as a microbial 'adaptation' to warming in the sense of microbes physiologically acclimating, rather than genetically evolving, to rising temperature[13,14].

Our ecosystem predictions, however, contrast with previous models in which microbial turnover (death rate) increases with temperature[4]. In our model, the microbial death rate has very little effect on the SOC ecosystem equilibrium, $C$, whereas Hagerty *et al.* (2014) found $C$ to increase with higher microbial turnover. The difference stems from the rate of enzyme production being resource dependent in our model, rather than constitutive and constant as in Allison *et al.* (2010) and Hagerty *et al.* (2014). Equilibrium $C$ is determined by the decomposition rate, which is controlled by the enzyme stock, $Z$. When exoenzyme production is constitutive, $Z$ is directly proportional to microbial biomass, $M$, and thus strongly influenced by the microbial death rate. When exoenzyme production is resource dependent, $Z$ becomes much more sensitive to the enzyme allocation parameters, $\varphi$, $\gamma_Z$ and $\gamma_M$. In our model, the temperature dependence of $d_M$ affects the ECOSresponse more through the dependence of $\varphi^*(T_0)$ on $d_M(T_0)$ (equation (S5)) than the dependence of $M$ on $d_M$.

**Supplementary Note 6: Evolutionary response of carbon-use efficiency (CUE)**

In our formalism, the microbial strategy of resource allocation to either microbial biomass synthesis or exoenzyme production follows a standard 'Y model' of resource allocation [15]. The exoenzyme production efficiency is denoted by $\gamma_Z$, and the microbial growth efficiency (MGE), by $\gamma_M$. Thus $(1 - \gamma_Z)$ measures the energetic cost of enzyme production and $(1 - \gamma_M)$ measures the energetic cost of microbial biomass synthesis; these costs are paid through microbial respiration returning $CO_2$ to the atmosphere (Fig. 1a). Larger costs may ensue from the production of enzymes needed to degrade more recalcitrant carbon compounds (smaller $\gamma_Z$); or the synthesis of more complex microbial structures (smaller $\gamma_M$).

Carbon-use efficiency (CUE) has received much attention in experimental studies, prompted by the contrasted effects of warming on respiration rates and microbial biomass[16–18]. Previous models have highlighted the effect of CUE on soil C stock[3], from local to global scale[19]. In our framework, CUE is given by

(S6) $$\text{CUE} = (1 - \varphi)\,\gamma_M + \varphi\,\gamma_Z$$

and thus appears as a compound parameter rather than a microbial trait[20]. Our model predicts that CUE changes as the microbial trait $\varphi$ evolves. Assuming constant MGE ($\gamma_M$), we find that natural selection always favors greater enzyme allocation fraction ($\varphi$) as temperature rises in the baseline scenario. From Supplementary Equation (6) we conclude that CUE decreases in microbes characterized by relatively costly exoenzymes and 'cheap' tissue ($\gamma_M < \gamma_Z$), and increases otherwise. Thus, even if MGE is kept constant with respect to temperature, microbial evolutionary adaptation to warming can have mixed effects on CUE.

**<u>Supplementary Note 7</u>: Influence of temperature dependence on SOC eco-evolutionary response**

If MGE decreases with temperature, the direction and magnitude of the EVO response to warming vary dramatically with the initial temperature $T_0$ (Fig. 3d, Supplementary Fig. 5d, h). The strong influence of $T_0$ on the ecosystem response of soil C to warming was noted by Li *et al.* (2014) for intermediate MGE temperature sensitivity ($m$ = - 0.008). The outstanding eco-evolutionary response (C sequestration) predicted by our model for steeper MGE temperature dependence ($m$ = - 0.014, Fig. 3d) are caused by the combination of $v_{max}^{U}(T)$ as an Arrhenius function with $\gamma_M(T)$ as a linear function (equations (2c) and (4) in Methods). While there is empirical support for this choice (reviewed in Todd-Brown *et al.* 2012[21]), both relationships remain uncertain and warrant further experimental investigation.

If the microbial turnover increases with temperature, warming selects for lower enzyme allocation fraction (Supplementary Fig. 5b, c), slowing down decomposition (Supplementary Fig. 5f, g), and possibly buffering the ecology-driven loss of soil C (Fig. 3b-c). Thus, when taking microbial evolution into account, the thermal sensitivity of microbial turnover does reduce C loss but never leads to the kind of C sequestration predicted by earlier purely ecosystem models[4].

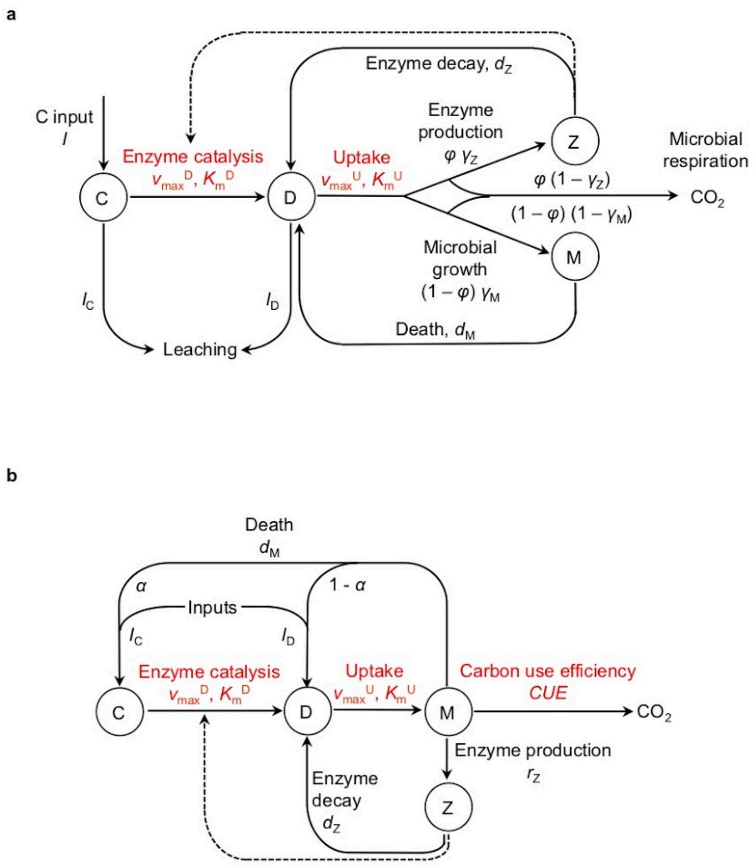


**Supplementary Figure 1: Structure of soil microbial decomposition models. (a)** Our model assumes dynamic allocation of assimilated carbon to enzyme production, whereas in **(b)** Allison *et al.* (2010)'s model, enzymes are produced at a constant rate.

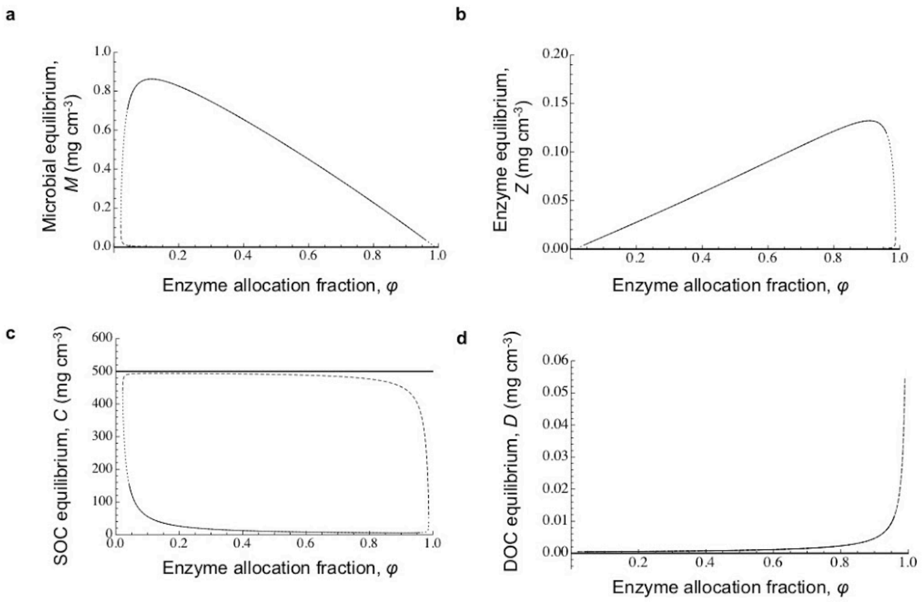


**Supplementary Figure 2: The three ecosystem equilibria.** Effect of enzyme allocation fraction, $\varphi$, on equilibrium existence and stability. *Thick line*, trivial stable equilibrium. *Dashed curve*, unstable equilibrium. *Plain curve*, non trivial stable equilibrium. *Dotted curve*, unstable equilibrium inside stable limit cycle. **(a)** Microbial biomass ($M$). **(b)** Exoenzyme concentration ($Z$). **(c)** SOC concentration ($C$). **(d)** DOC concentration ($D$). All state variables are measured in unit mass of carbon. Parameters are set to their default values (Supplementary Table 1).

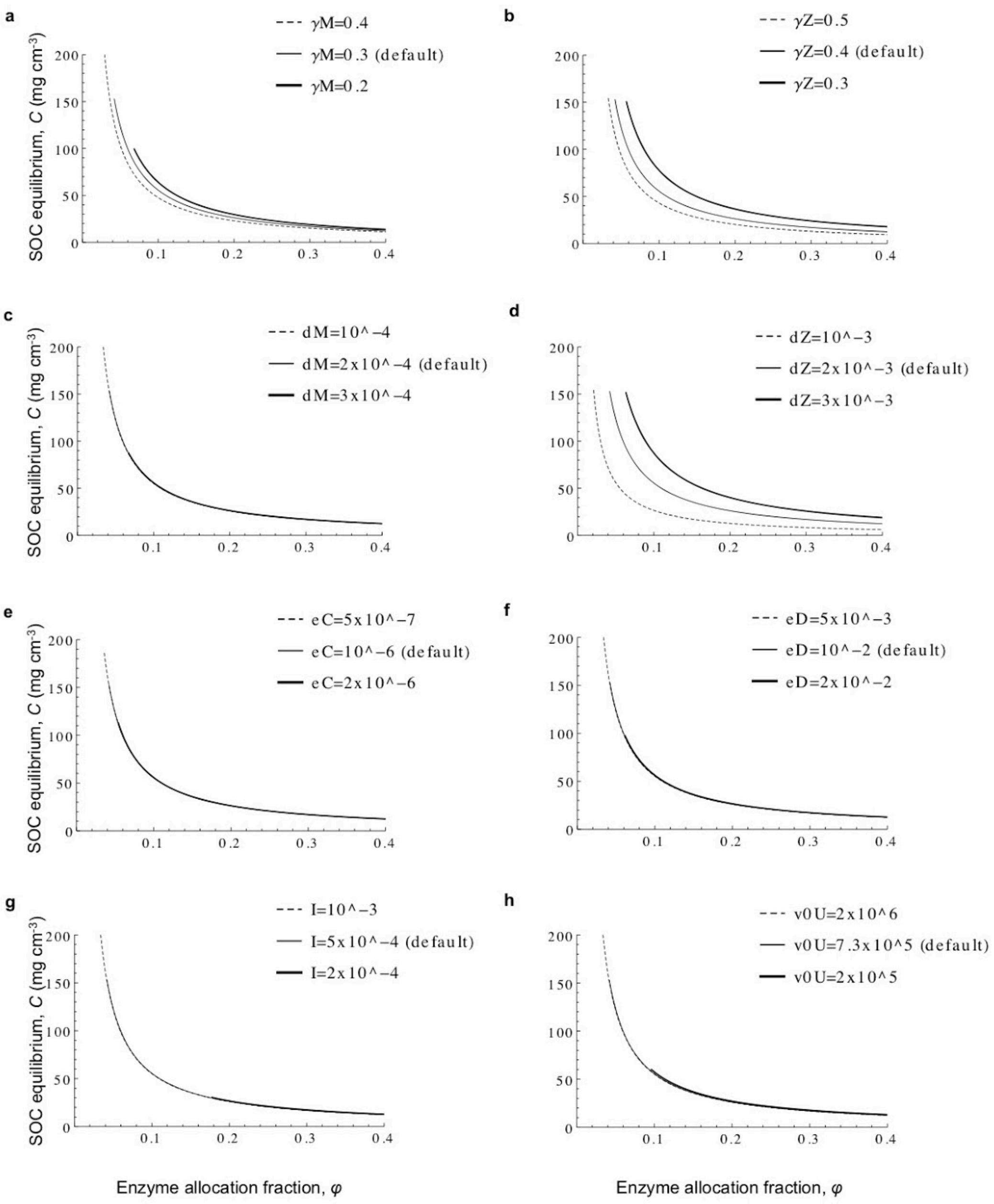
a
γM=0.4
γM=0.3 (default)
γM=0.2
b
γZ=0.5
γZ=0.4 (default)
γZ=0.3
c
dM=10^−4
dM=2x10^−4 (default)
dM=3x10^−4
d
dZ=10^−3
dZ=2x10^−3 (default)
dZ=3x10^−3
e
eC=5x10^−7
eC=10^−6 (default)
eC=2x10^−6
f
eD=5x10^−3
eD=10^−2 (default)
eD=2x10^−2
g
I=10^−3
I=5x10^−4 (default)
I=2x10^−4
h
v0U=2x10^6
v0U=7.3x10^5 (default)
v0U=2x10^5
SOC equilibrium, C (mg cm-3)
Enzyme allocation fraction, φ
0
50
100
150
200
0.1
0.2
0.3
0.4

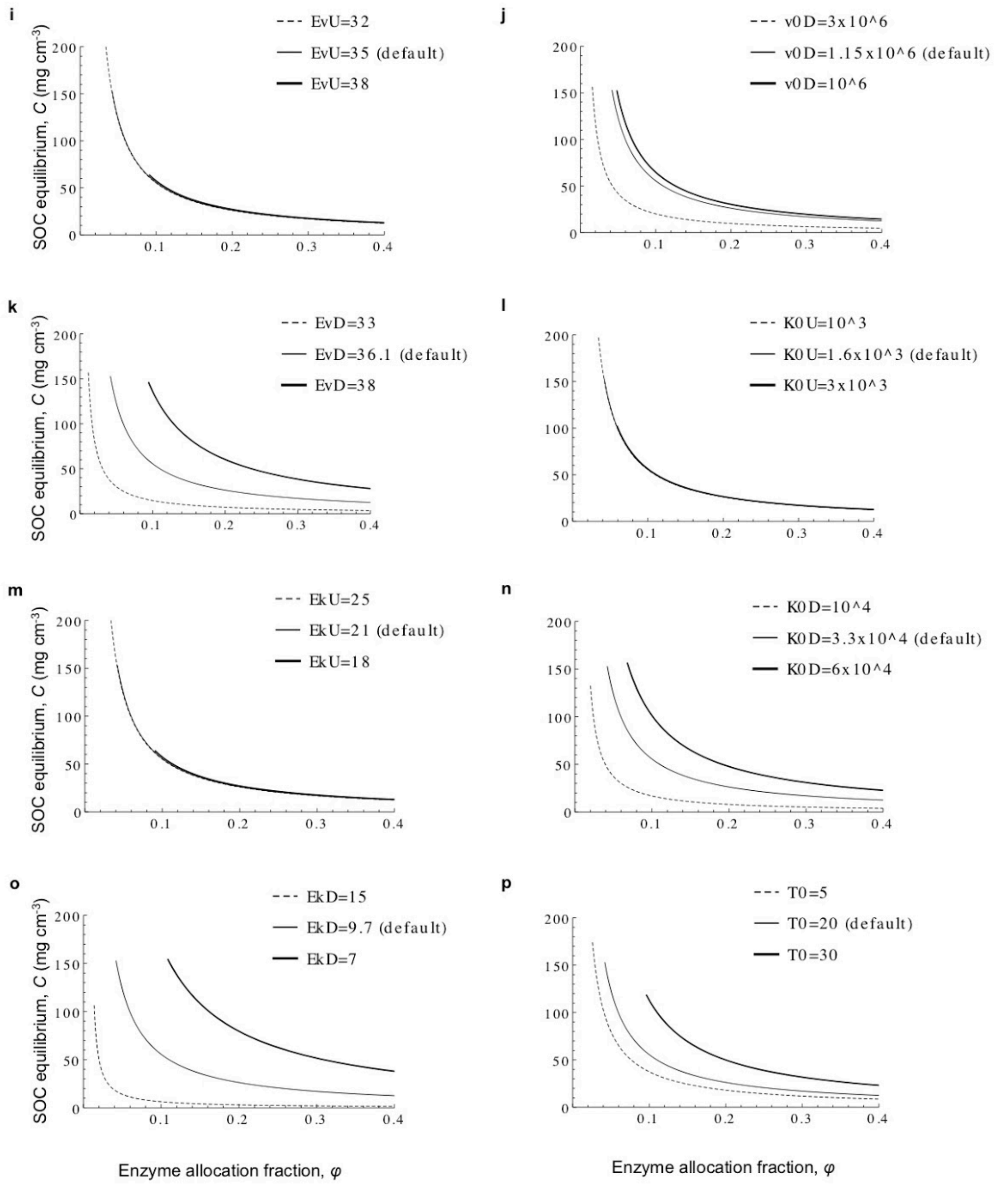


<u>Supplementary Figure 3</u>: **Parameter influence on the dependence of equilibrium *C* on enzyme allocation fraction, *φ*. (a)** Microbial growth efficiency, $\gamma_M$. **(b)** Enzyme production efficiency, $\gamma_Z$. **(c)** Microbial mortality rate, $d_M$. **(d)** Enzyme deactivation rate, $d_Z$. **(e)** SOC leaching rate, $e_C$. **(f)** DOC leaching rate, $e_D$. **(g)** SOC input (litter), *I*. **(h)** Arrhenius coefficient of uptake rate, $v_0^U$. **(i)** Activation energy of uptake rate, $E_v^U$. **(j)** Arrhenius coefficient of decomposition rate, $v_0^D$. **(k)** Activation energy of decomposition rate, $E_v^D$. **(l)** Arrhenius

coefficient of uptake half-saturation constant, $K_0^{U}$. **(m)** Activation energy of uptake half-saturation constant, $E_K^{U}$. **(n)** Arrhenius coefficient of decomposition half-saturation constant, $K_0^{D}$. **(o)** Activation energy of decomposition half-saturation constant, $E_K^{D}$. **(p)** Initial temperature, $T_0$. Two parameter values (thick and dashed curves) are tested around the default value (plain curve). Other parameters are set to their default values (Supplementary Table 1).

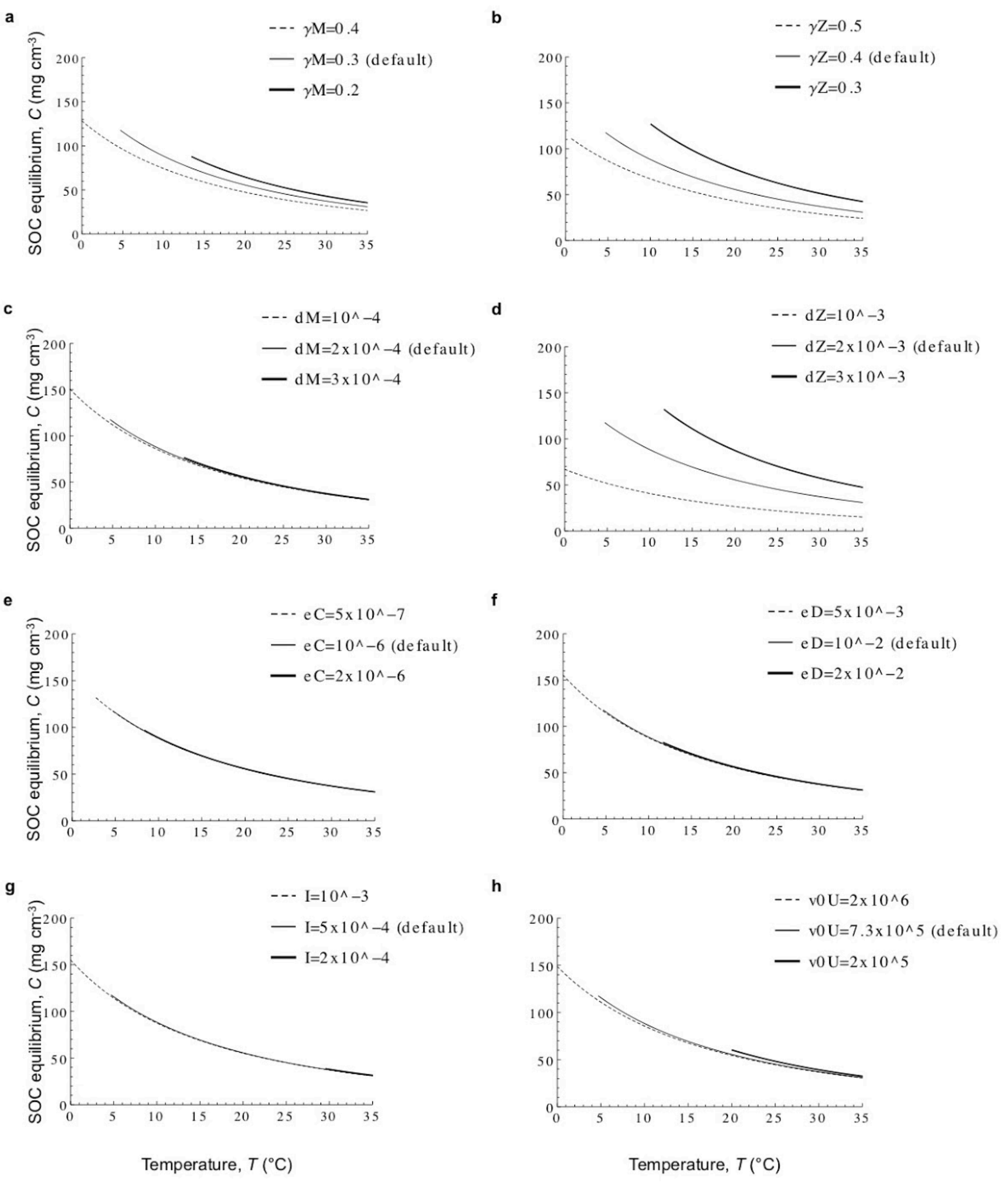

a
γM=0.4
γM=0.3 (default)
γM=0.2
b
γZ=0.5
γZ=0.4 (default)
γZ=0.3
c
dM=10^−4
dM=2x10^−4 (default)
dM=3x10^−4
d
dZ=10^−3
dZ=2x10^−3 (default)
dZ=3x10^−3
e
eC=5x10^−7
eC=10^−6 (default)
eC=2x10^−6
f
eD=5x10^−3
eD=10^−2 (default)
eD=2x10^−2
g
I=10^−3
I=5x10^−4 (default)
I=2x10^−4
h
v0U=2x10^6
v0U=7.3x10^5 (default)
v0U=2x10^5
SOC equilibrium, C (mg cm⁻³)
Temperature, T (°C)

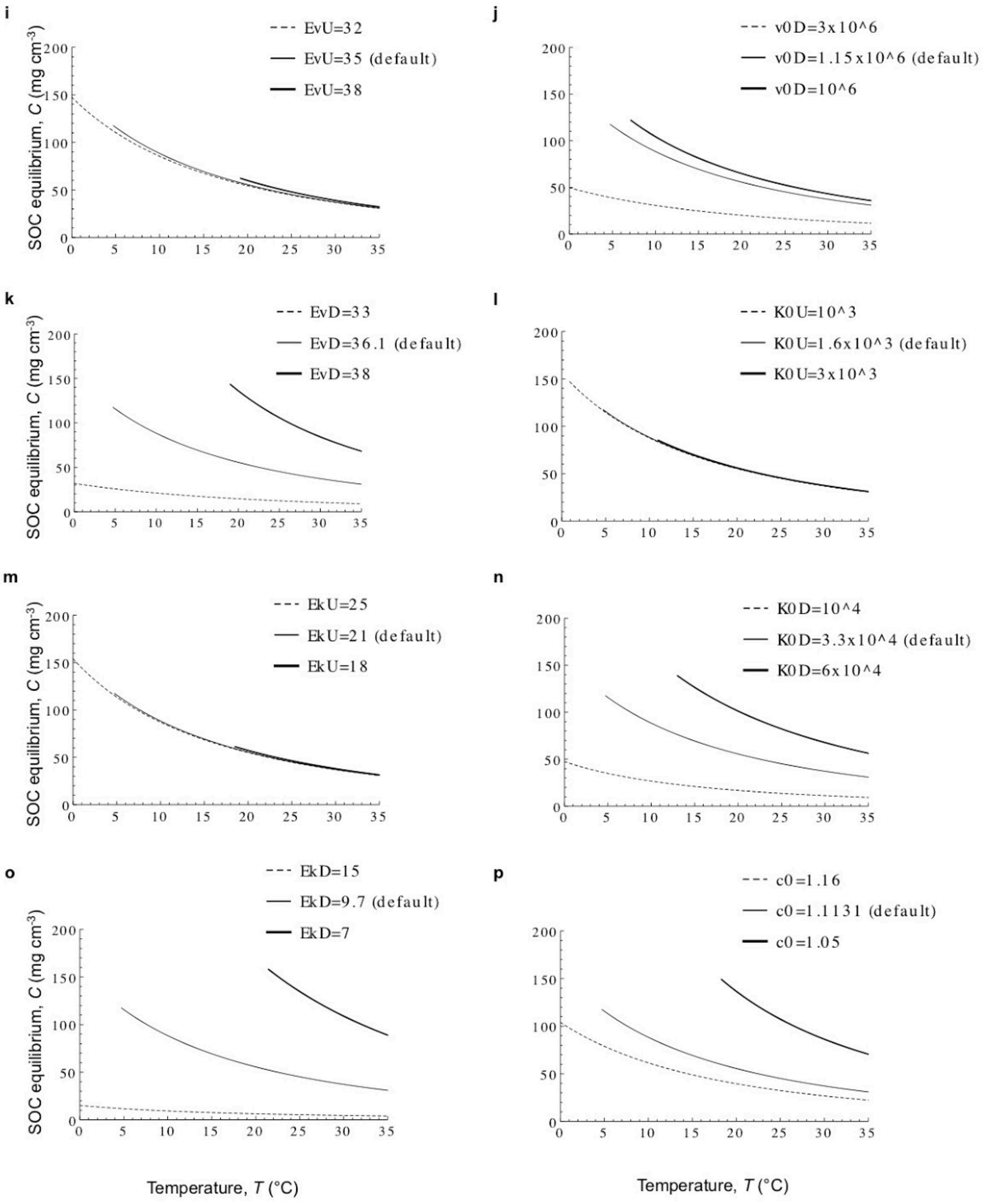


**Supplementary Figure 4: Parameter influence on the dependence of equilibrium *C* on temperature, *T*.** Baseline 'kinetics-only' scenario of temperature dependence. **(a)** Microbial growth efficiency, $\gamma_M$. **(b)** Enzyme production efficiency, $\gamma_Z$. **(c)** Microbial mortality rate, $d_M$. **(d)** Enzyme deactivation rate, $d_Z$. **(e)** SOC leaching rate, $e_C$. **(f)** DOC leaching rate, $e_D$. **(g)** SOC input (litter), $I$. **(h)** Arrhenius coefficient of uptake rate, $v_0^U$. **(i)** Activation energy of uptake rate, $E_v^U$. **(j)** Arrhenius coefficient of decomposition rate, $v_0^D$. **(k)** Activation energy of decomposition rate,

$E_v^D$. **(l)** Arrhenius coefficient of uptake half-saturation constant, $K_0^U$. **(m)** Activation energy of uptake half-saturation constant, $E_K^U$. **(n)** Arrhenius coefficient of decomposition half-saturation constant, $K_0^D$. **(o)** Activation energy of decomposition half-saturation constant, $E_K^D$. **(p)** Competition asymmetry, $c_0$. Two parameter values (thick and dashed curves) are tested around the default value (plain curve). For each parameter value, enzyme allocation fraction, $\varphi$, is equal to the adapted value, $\varphi^*$, at $T$ = 20°C. Other parameters are set to their default values (Supplementary Table 1).

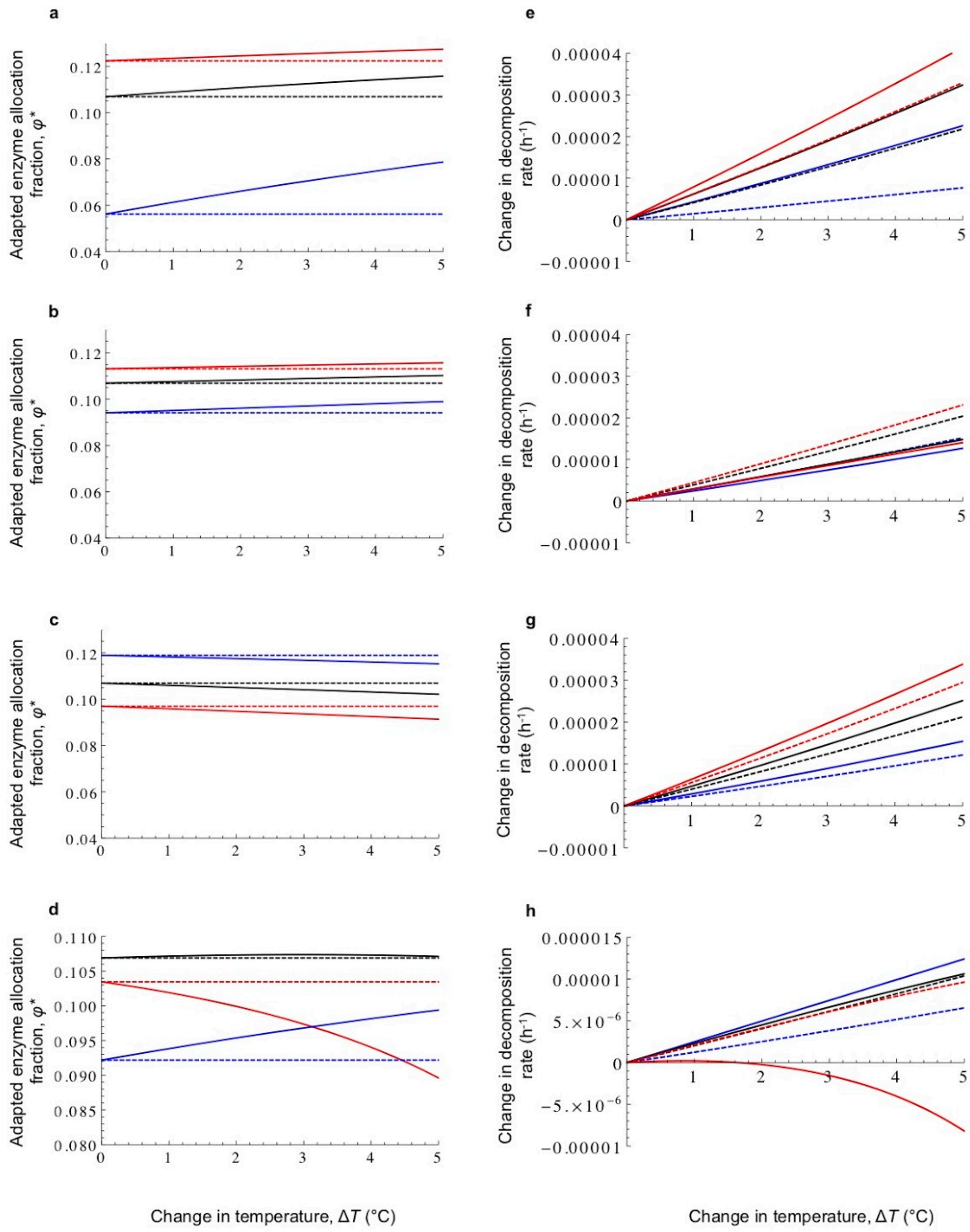


**<u>Supplementary Figure 5</u>: Ecosystem and eco-evolutionary responses of the adaptive enzyme allocation fraction, $\varphi^*$ (a-d), and decomposition rate (e-h) to warming for three scenarios of temperature dependence.** Ecosystem response (without evolution, dashed curves) and eco-evolutionary response (with evolution, plain curves) are plotted as a function of the increase in temperature, up to + 5°C. *Blue curves*, initial temperature $T_0$ = 5°C. *Black curves*, $T_0$ = $T_{ref}$ = 20°C. *Red curves*, $T_0$ = 30°C. **(a, e)** Baseline 'kinetics only' scenario of temperature dependence. **(b, f)** Temperature-dependent microbial turnover, with $E_{dM} = 25 < E_v^U$. **(c, g)** Temperature-dependent microbial turnover, with $E_{dM} = 55 > E_v^U$. **(d, h)** Temperature-dependent MGE, with $m = -0.014$. Parameters are set to their default values (Supplementary Table 1), except $I = 5\ 10^{-3}$, $v_0^U = 10^5$, $E_v^U = 38$, $c_0 = 1.17$.

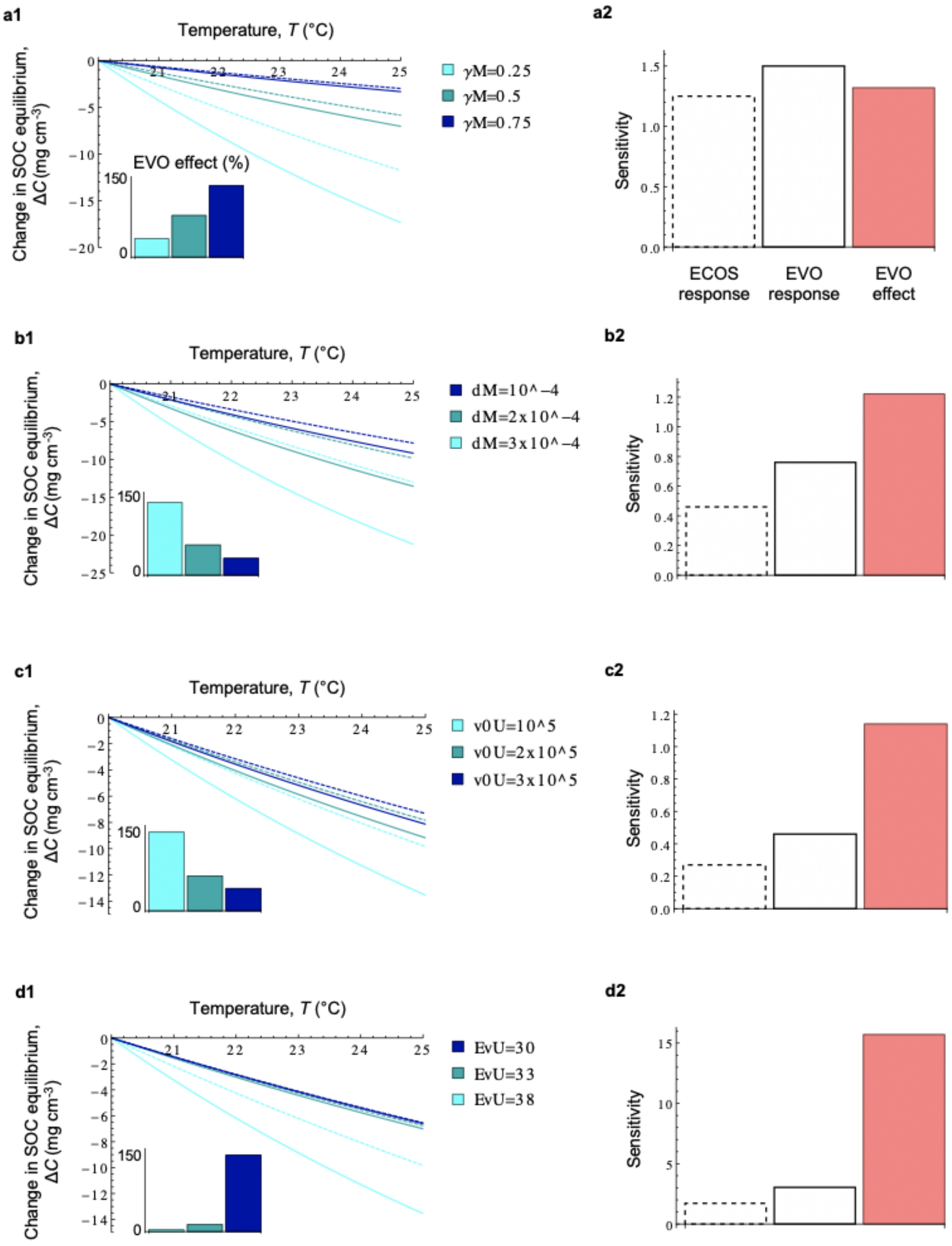
a1
Temperature, T (°C)
Change in SOC equilibrium, ΔC (mg cm⁻³)
EVO effect (%)
γM=0.25
γM=0.5
γM=0.75
a2
Sensitivity
ECOS response
EVO response
EVO effect
b1
dM=10^−4
dM=2x10^−4
dM=3x10^−4
b2
c1
v0U=10^5
v0U=2x10^5
v0U=3x10^5
c2
d1
EvU=30
EvU=33
EvU=38
d2

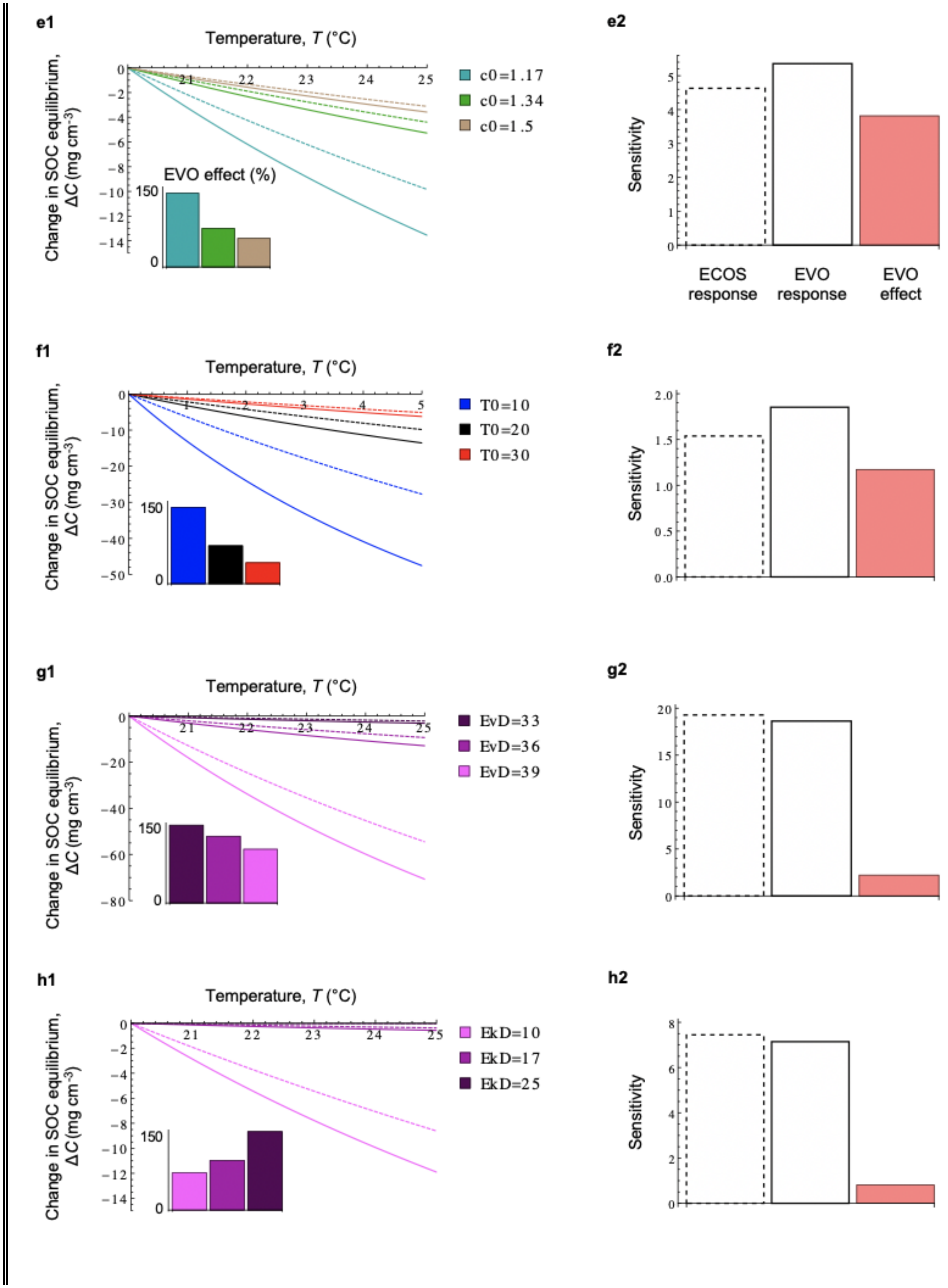
e1
Temperature, T (°C)
Change in SOC equilibrium, ΔC (mg cm-3)
0
−2
−4
−6
−8
−10
−12
−14
21
22
23
24
25
c0=1.17
c0=1.34
c0=1.5
EVO effect (%)
150
0
e2
Sensitivity
0
1
2
3
4
5
ECOS response
EVO response
EVO effect
f1
Temperature, T (°C)
Change in SOC equilibrium, ΔC (mg cm-3)
0
−10
−20
−30
−40
−50
1
2
3
4
5
T0=10
T0=20
T0=30
150
0
f2
Sensitivity
0.0
0.5
1.0
1.5
2.0
g1
Temperature, T (°C)
Change in SOC equilibrium, ΔC (mg cm-3)
0
−20
−40
−60
−80
21
22
23
24
25
EvD=33
EvD=36
EvD=39
150
0
g2
Sensitivity
0
5
10
15
20
h1
Temperature, T (°C)
Change in SOC equilibrium, ΔC (mg cm-3)
0
−2
−4
−6
−8
−10
−12
−14
21
22
23
24
25
EkD=10
EkD=17
EkD=25
150
0
h2
Sensitivity
0
2
4
6
8

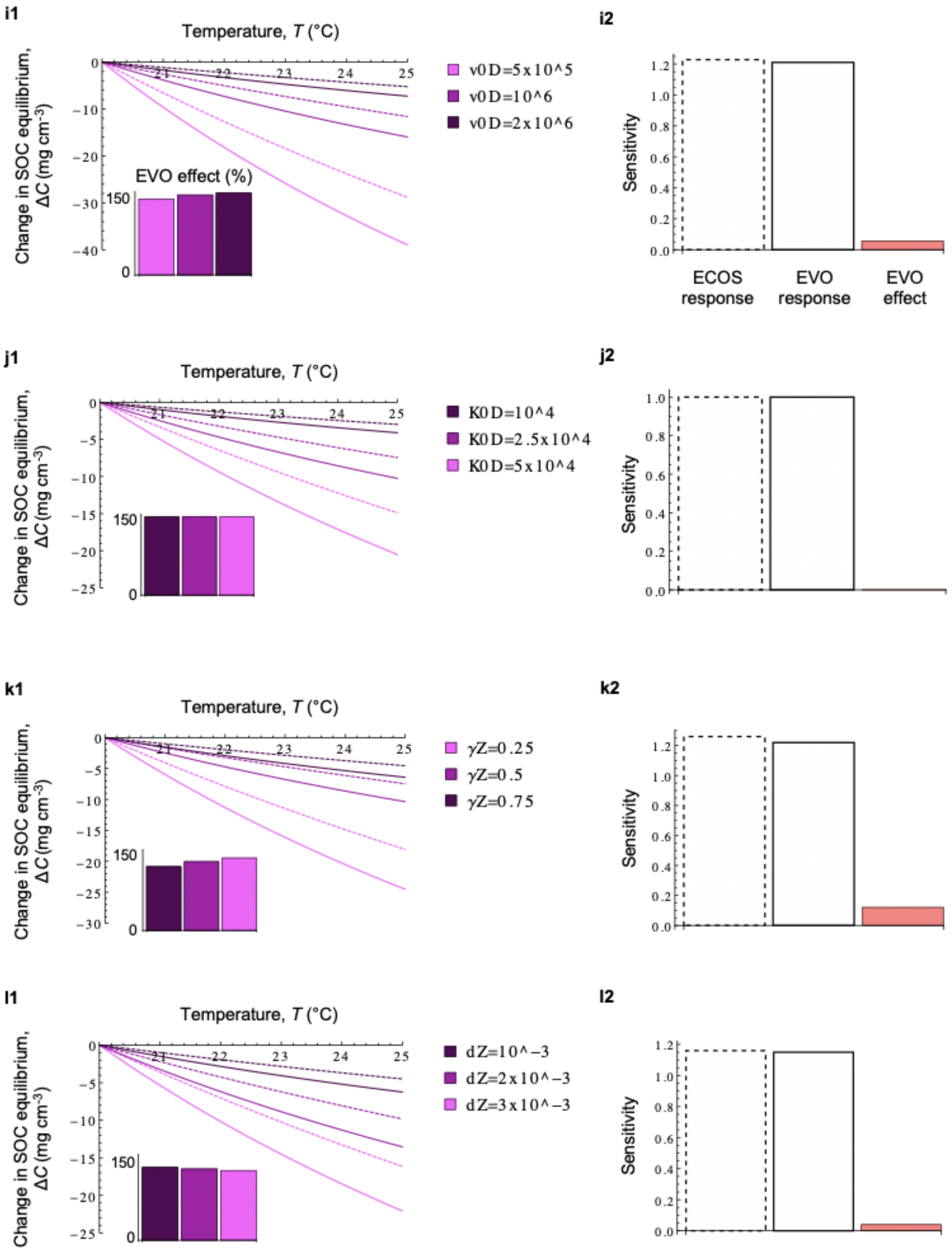
i1
Temperature, T (°C)
Change in SOC equilibrium, ΔC (mg cm⁻³)
EVO effect (%)
v0D=5x10^5
v0D=10^6
v0D=2x10^6
i2
Sensitivity
ECOS response
EVO response
EVO effect
j1
Temperature, T (°C)
Change in SOC equilibrium, ΔC (mg cm⁻³)
K0D=10^4
K0D=2.5x10^4
K0D=5x10^4
j2
Sensitivity
k1
Temperature, T (°C)
Change in SOC equilibrium, ΔC (mg cm⁻³)
γZ=0.25
γZ=0.5
γZ=0.75
k2
Sensitivity
l1
Temperature, T (°C)
Change in SOC equilibrium, ΔC (mg cm⁻³)
dZ=10^−3
dZ=2x10^−3
dZ=3x10^−3
l2
Sensitivity

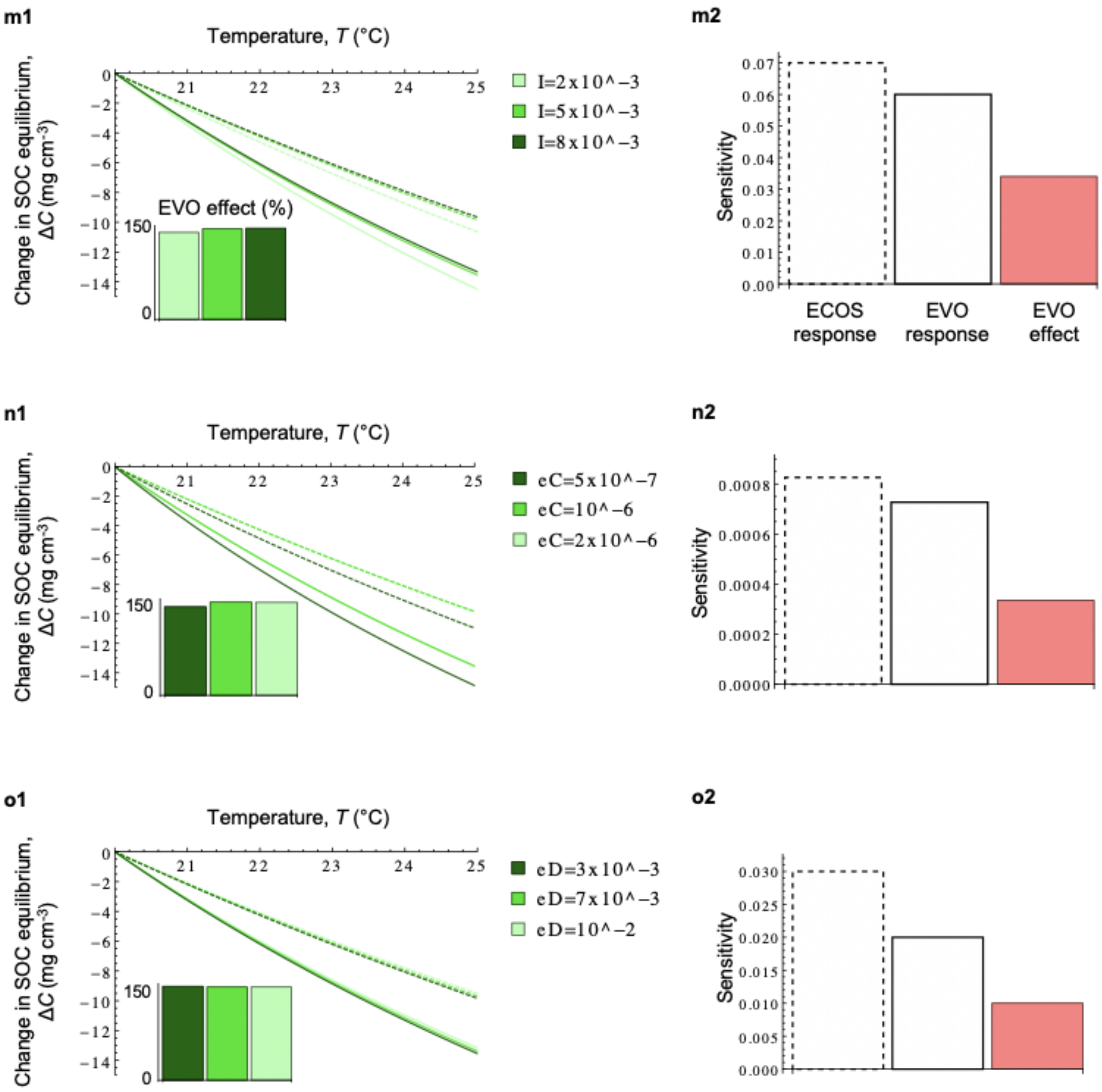


**<u>Supplementary Figure 6</u>: Parameter influence on the ECOS response, EVO response and EVO effect.** Baseline 'kinetics-only' scenario of temperature dependence. *Left column*, ECOSresponse (dashed curves), EVO response (plain curves), and EVO effect (insets), for three parameters values. *Right column*, Sensitivity of ECOS(dashed), EVO (plain) and EVO (pink) to each parameter. Sensitivity is calculated with the minimal and maximal values tested for each parameter and is expressed in absolute value. Because we lack information about realistic range for most parameters, we show relative rather than absolute sensitivities (and the scale for the y-axis is different for each parameter). **(a1-a2)** Microbial growth efficiency, $\gamma_M$. **(b1-b2)** Microbial mortality rate, $d_M$. **(c1-c2)** Arrhenius coefficient of uptake rate, $v_0^U$. **(d1-d2)** Activation energy of uptake rate, $E_v^U$. **(e1-e2)** Competition asymmetry, $c_0$. **(f1-f2)** Initial temperature, $T_0$. **(g1-g2)** Activation energy of decomposition rate, $E_v^D$. **(h1-h2)** Activation energy of decomposition half-saturation constant, $E_K^D$. **(i1-i2)** Arrhenius coefficient of decomposition rate, $v_0^D$. **(j1-j2)** Arrhenius coefficient of decomposition half-saturation constant, $K_0^D$. **(k1-k2)** Enzyme production efficiency, $\gamma_Z$. **(l1-l2)** Enzyme deactivation rate, $d_Z$. **(m1-m2)** SOC input (litter), $I$. **(n1-n2)** SOC leaching rate, $e_C$. **(o1-o2)** DOC leaching rate, $e_D$. Parameters are set to their default values (Supplementary Table 1) except $I = 5\ 10^{-3}$, $v_0^U = 10^5$, $E_v^U = 38$, $c_0 = 1.17$. Darker blue indicates higher microbial performance, darker purple indicates higher enzyme performance, darker green indicates higher soil carbon retention (higher $I$, lower $e_C$, lower $e_D$).

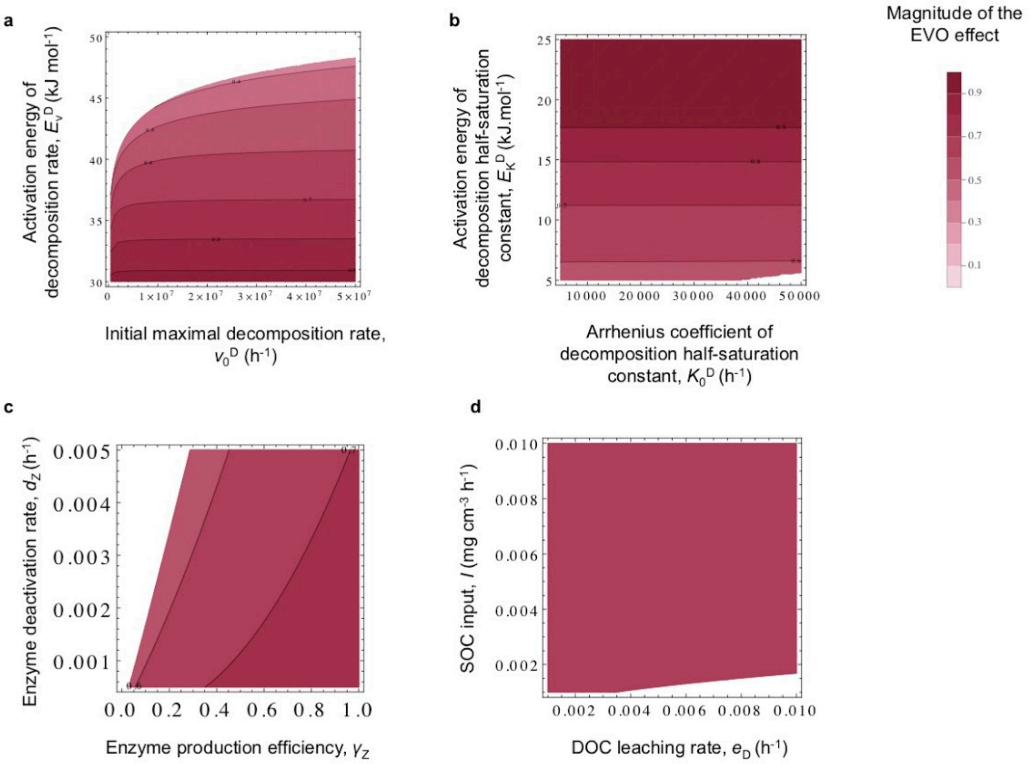


**Supplementary Figure 7: Sensitivity analysis of the EVO effect for the baseline 'kinetics-only' scenario of temperature dependence. (a)** Sensitivity to Arrhenius coefficient of decomposition rate, $v_0^D$, and activation energy of decomposition rate, $E_v^D$. **(b)** Sensitivity to Arrhenius coefficient of decomposition half-saturation constant, $K_0^D$, and activation energy of decomposition half-saturation constant, $E_K^D$. **(c)** Sensitivity to enzyme production efficiency, $\gamma_Z$, and enzyme deactivation rate, $d_Z$. **(d)** Sensitivity to DOC leaching rate, $e_D$, and SOC input (litter), $I$. Parameters are set to their default values (Supplementary Table 1) except $I = 5\ 10^{-3}$, $v_0^U = 10^5$, $E_v^U = 38$ (point B2). The effect of $e_C$ (not shown) is identical to the effect of $e_D$.

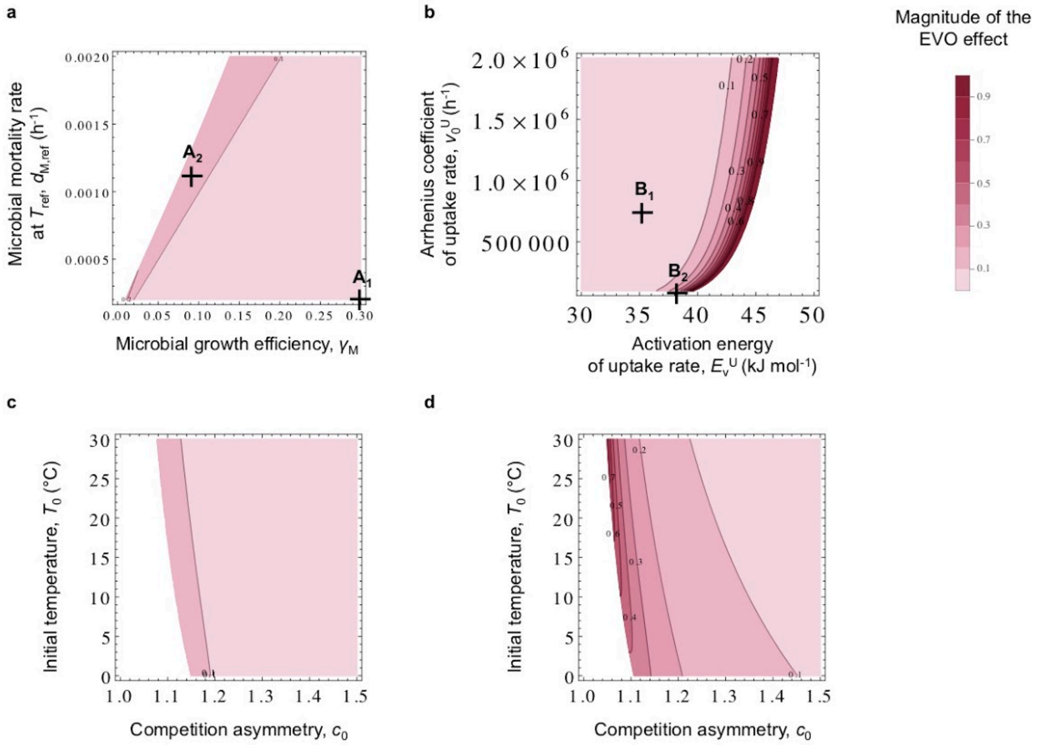
a
b
c
d
Magnitude of the EVO effect
Microbial mortality rate at $T_{ref}$, $d_{M,ref}$ ($h^{-1}$)
Microbial growth efficiency, $\gamma_M$
$A_1$
$A_2$
Arrhenius coefficient of uptake rate, $v_0^U$ ($h^{-1}$)
Activation energy of uptake rate, $E_v^U$ (kJ $mol^{-1}$)
$B_1$
$B_2$
Initial temperature, $T_0$ (°C)
Competition asymmetry, $c_0$

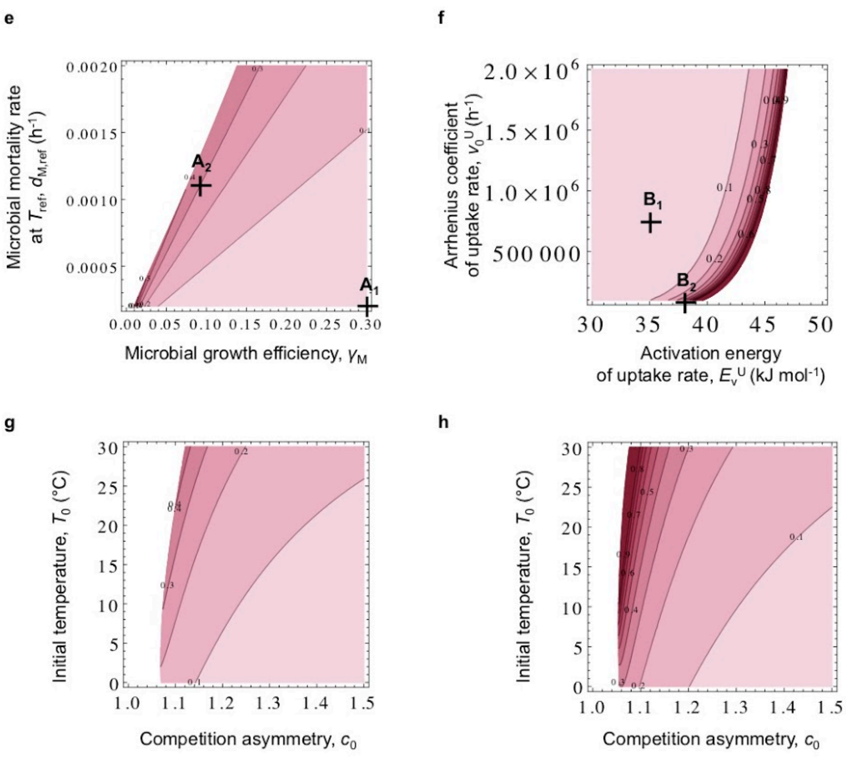

e
f
g
h
A2
A1
B1
B2
Microbial mortality rate at $T_{ref}$, $d_{M,ref}$ (h$^{-1}$)
Microbial growth efficiency, $\gamma_M$
Arrhenius coefficient of uptake rate, $v_0^U$ (h$^{-1}$)
Activation energy of uptake rate, $E_v^U$ (kJ mol$^{-1}$)
Initial temperature, $T_0$ (°C)
Competition asymmetry, $c_0$
Initial temperature, $T_0$ (°C)
Competition asymmetry, $c_0$

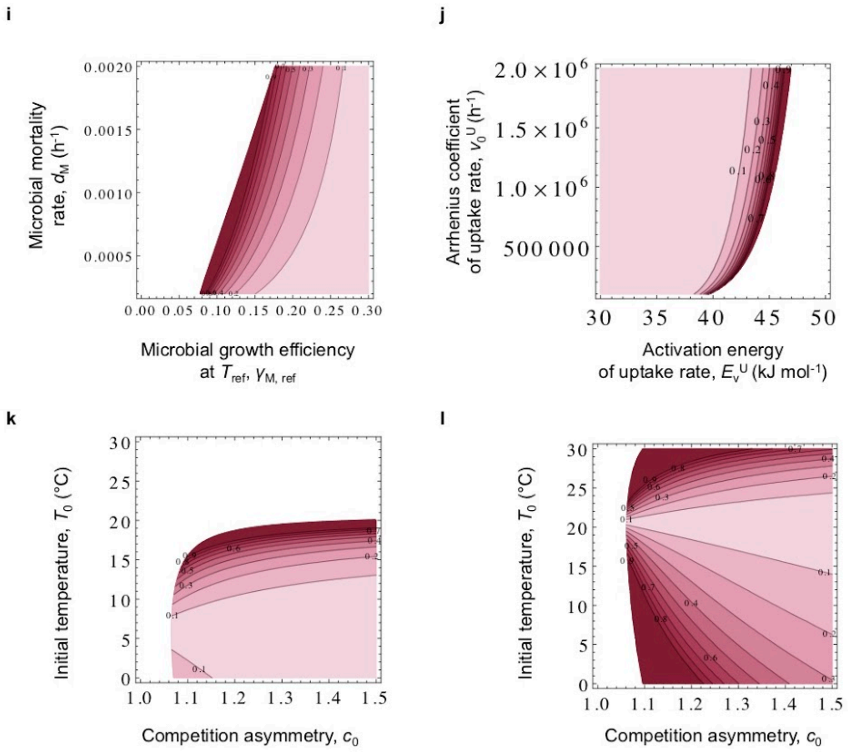


**<u>Supplementary Figure 8</u>: Sensitivity analysis of the EVO effect for the microbial turnover and MGE scenarios of temperature dependence. (a-d)** Temperature-dependent microbial turnover, $E_{dM} = 25$. **(e-h)** Temperature-dependent microbial turnover, $E_{dM} = 55$. **(i-l)** Temperature-dependent MGE, $m = -\ 0.014$. Parameters are set to their default values (Supplementary Table 1) except $I = 5\ 10^{-3}$. A1 = B1 = default values. A2 = default values except for $\gamma_M$ and $d_M$. B2 = default values except for $v_0^U$ and $E_v^U$. **(a, e, i)** Sensitivity to microbial growth efficiency, $\gamma_M$, and microbial mortality rate, $d_M$. **(b, f, j)** Sensitivity to activation energy of uptake rate, $E_v^U$, and Arrhenius coefficient of uptake rate, $v_0^U$. **(c, d, g, h, k, l)** Sensitivity to competition asymmetry, $c_0$, and initial temperature, $T_0$. Panels **(c, j, k)** (resp. **d, h, l**) show sensitivities of the EVO effect at point A2 (resp. B2) for the corresponding scenario of temperature dependence.

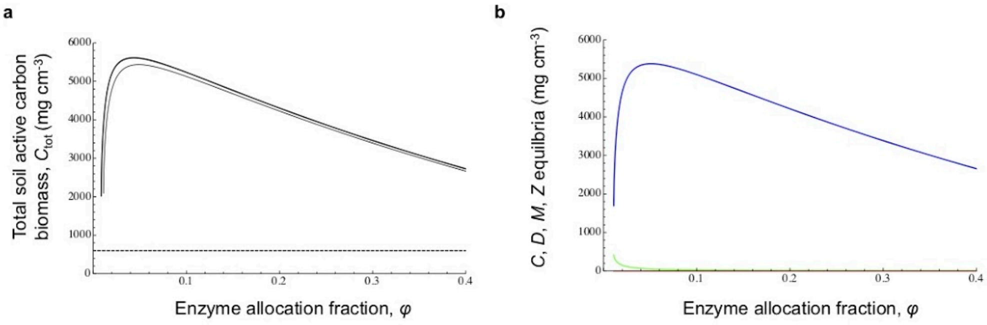


**<u>Supplementary Figure 9</u>: Effect of enzyme allocation fraction on total soil active carbon and its four components when microbial biomass forms the main part of active soil carbon.** **(a)** Total soil active carbon at 20°C (plain) and 25°C (bold) as a function of enzyme allocation fraction, $\varphi$. Note that the SOC equilibrium in the absence of microbes (dashed line) is temperature independent and always lower than the total soil active carbon. **(b)** Effect of enzyme allocation fraction, $\varphi$, on the four components components of total soil active carbon at 20°C: *M* (blue), *Z* (pink), *C* (green), *D* (orange). Parameters are set to their default values (Supplementary Table 1) except $I = 3\ 10^{-3}$, $v_0^U = 4\ 10^3$, $E_v^U = 46$, $d_M = 10^{-6}$, $e_C = 5\ 10^{-6}$ and $\gamma_M = 0.7$.

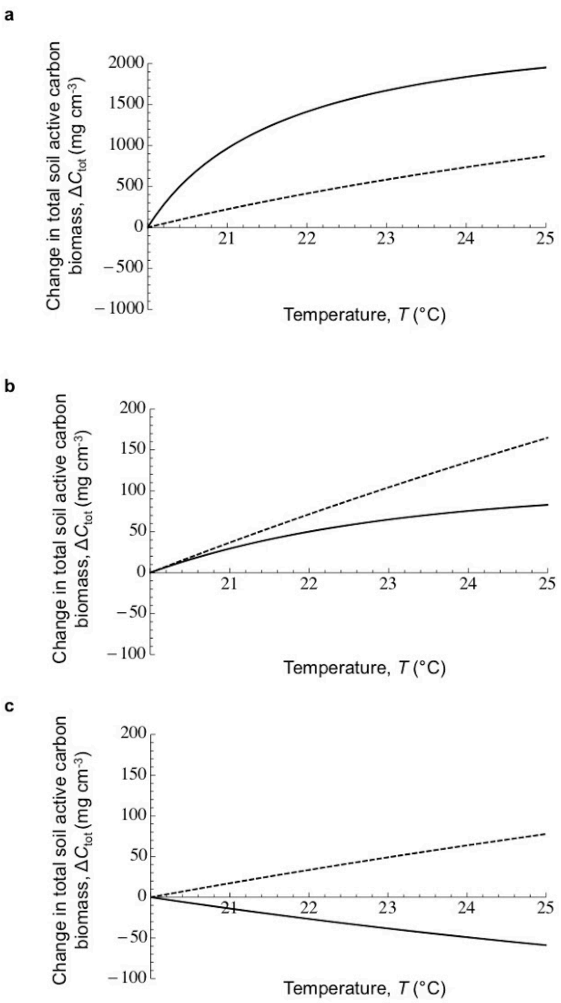


**<u>Supplementary Figure 10</u>: ECOSand EVO responses when microbial biomass forms the main part of active soil carbon for different local competitive advantage to enzyme producers (competition asymmetry). (a)** Small advantage ($c_0$ = 1.075). **(b)** Intermediate advantage ($c_0$ = 1.12). **(c)** Large advantage ($c_0$ = 1.35). Parameters are set to their default values (Supplementary Table 1) except $I = 3\ 10^{-3}$, $v_0^{U} = 4\ 10^{3}$, $E_v^{U} = 46$, $d_M = 10^{-6}$, $e_C = 5\ 10^{-6}$ and $\gamma_M$ = 0.7.

| Parameter | Unit | Description | Default value |
|---|---|---|---|
| $T_0$, $T_{ref}$ | °C | initial temperature | 20 |
| $\varphi$ | | enzyme allocation fraction | 0.1 |
| $\gamma_M$, $\gamma_{M,ref}$ | | microbial growth efficiency | 0.3, 0.31 |
| $\gamma_Z$ | | enzyme production efficiency | 0.4 |
| $d_M$, $d_{M,ref}$ | $h^{-1}$ | microbial mortality rate | $2\ 10^{-4}$ |
| $d_Z$ | $h^{-1}$ | enzyme deactivation rate | $2\ 10^{-3}$ |
| $v_0^U$ | mg D $cm^{-3}$ (mg M $cm^{-3}$)$^{-1}$ $h^{-1}$ | Arrhenius coefficient of uptake rate | $7.3\ 10^{5}$ |
| $v_0^D$ | mg C $cm^{-3}$ (mg Z $cm^{-3}$)$^{-1}$ $h^{-1}$ | Arrhenius coefficient of decomposition rate | $1.15\ 10^{6}$ |
| $K_0^U$ | mg D $cm^{-3}$ | Arrhenius coefficient of uptake half-saturation constant | $1.6\ 10^{3}$ |
| $K_0^D$ | mg C $cm^{-3}$ | Arrhenius coefficient of decomposition half-saturation constant | $3.3\ 10^{4}$ |
| $E_v^U$ | kJ $mol^{-1}$ | activation energy of uptake rate | 35 |
| $E_v^D$ | kJ $mol^{-1}$ | activation energy of decomposition rate | 36.1 |
| $E_K^U$ | kJ $mol^{-1}$ | activation energy of uptake half-saturation constant | 21 |
| $E_K^D$ | kJ $mol^{-1}$ | activation energy of decomposition half-saturation constant | 9.7 |
| $E_{dM}$ | kJ $mol^{-1}$ | activation energy of microbial turnover | 0, 25, 55 |
| $I$ | mg C $cm^{-3}$ $h^{-1}$ | SOC input (litter) | $5\ 10^{-4}$ |
| $e_C$ | $h^{-1}$ | SOC leaching rate | $10^{-6}$ |
| $e_D$ | $h^{-1}$ | DOC leaching rate | $10^{-2}$ |
| $c_0$ | | local competitive advantage to stronger exoenzyme producers (competition asymmetry) | 1.1131 |
| $\Delta T$ | °C | warming treatment | 5 |

**Supplementary Table 1: Default parameter values.**

| Parameter | Low value | High value | Sensitivity (absolute value) | | | | Qualitative effect on $\varphi_{min}$, $\varphi_{max}$ and equilibrium $C$ |
|---|---|---|---|---|---|---|---|
| | | | C | D | M | Z | |
| $T$ | 4.9 | 490 | 1.62 | 0.83 | 0.06 | 0.06 | Lower $\varphi_{min}$, higher $\varphi_{max}$, $C$ decreases especially for low values of $T$. |
| $\varphi$ | 0.19 | 0.95 | 0.97 | 1.75 | 1.75 | 0.98 | $C$ decreases especially for low values of $\varphi$. |
| $\gamma_M$ | 0.19 | 0.95 | 1.32 | 1 | 2.34 | 1.34 | Lower $\varphi_{min}$, higher $\varphi_{max}$, $C$ decreases especially for very high values of $\gamma_M$. |
| $\gamma_Z$ | 0.19 | 0.95 | 1.18 | 0 | 0.24 | 1.24 | Lower $\varphi_{min}$, higher $\varphi_{max}$, $C$ decreases strongly. |
| $d_M$ | $3.8\ 10^{-6}$ | $3.8\ 10^{-4}$ | 0.005 | 1 | 1.01 | 0.005 | Higher $\varphi_{min}$, lower $\varphi_{max}$, $C$ unchanged. |
| $d_Z$ | $4.6\ 10^{-5}$ | $4.6\ 10^{-3}$ | 1.05 | 0 | 0.07 | 1.07 | Higher $\varphi_{min}$, lower $\varphi_{max}$, $C$ increases strongly. |
| $v_0^U$ | $2.1\ 10^{5}$ | $2.1\ 10^{7}$ | 0.01 | 1 | 0.01 | 0.01 | Lower $\varphi_{min}$, higher $\varphi_{max}$, $C$ unchanged. |
| $v_0^D$ | $5.1\ 10^{5}$ | $5.1\ 10^{7}$ | 1.05 | 0 | 0.07 | 0.07 | Lower $\varphi_{min}$, higher $\varphi_{max}$, $C$ decreases strongly. |
| $K_0^U$ | 60 | 6000 | 0.01 | 1 | 0.01 | 0.01 | Higher $\varphi_{min}$, lower $\varphi_{max}$, $C$ unchanged. |
| $K_0^D$ | 900 | $9\ 10^{4}$ | 1 | 0 | 0.08 | 0.08 | Higher $\varphi_{min}$, lower $\varphi_{max}$, $C$ increases strongly. |
| $E_v^U$ | 0.38 | 38 | 0.01 | 3.35 | 0.01 | 0.01 | Higher $\varphi_{min}$, lower $\varphi_{max}$, $C$ unchanged. |
| $E_v^D$ | 0.38 | 38 | 3.4 | 0 | 0.07 | 0.07 | Higher $\varphi_{min}$, lower $\varphi_{max}$, $C$ increases strongly. |
| $E_K^U$ | 18 | 1800 | 0.01 | 159 | 0.01 | 0.01 | Lower $\varphi_{min}$, higher $\varphi_{max}$, $C$ unchanged. |
| $E_K^D$ | 7.3 | 730 | 7.58 | 0 | 0.08 | 0.08 | Lower $\varphi_{min}$, higher $\varphi_{max}$, $C$ decreases strongly. |
| $I$ | $2.8\ 10^{-4}$ | $2.8\ 10^{-2}$ | 0.005 | 0 | 1.05 | 1.05 | Lower $\varphi_{min}$, higher $\varphi_{max}$, $C$ unchanged. |

| | | | | | | | |
|---|---|---|---|---|---|---|---|
| $e_C$ | $5\ 10^{-8}$ | $5\ 10^{-6}$ | 0.003 | 0 | 0.18 | 0.18 | Higher $\varphi_{min}$, lower $\varphi_{max}$, $C$ unchanged. |
| $e_D$ | $3.8\ 10^{-4}$ | $3.8\ 10^{-2}$ | 0.01 | 0 | 0.01 | 0.01 | Higher $\varphi_{min}$, lower $\varphi_{max}$, $C$ unchanged. |
| $c_0$ | 1.1 | 25 | 0.75 | 1 | 0.99 | 0.76 | $C$ decreases especially for low values of $c_0$. |

**<u>Supplementary Table 2</u>: Sensitivity analysis of the ecosystem model.** The table reports the absolute value of sensitivities.

| Parameter | Location | | | | |
|---|---|---|---|---|---|
| | Alaska (AK) | Maine (ME) | West Virginia (WV) | California (CA) | Costa Rica (CR) |
| $T_0$ (°C) | 0 | 5 | 9 | 17 | 26 |
| $v_0^D$ ($h^{-1}$) | $7.7\ 10^{7}$ | $7.73\ 10^{9}$ | $1.35\ 10^{8}$ | $1.15\ 10^{6}$ | $1.23\ 10^{8}$ |
| $E_v^D$ (kJ $mol^{-1}$) | 43.7 | 50.6 | 47.2 | 36.1 | 48.9 |
| $K_0^D$ (mg $cm^{-3}$) | $2.79\ 10^{7}$ | $2.1\ 10^{7}$ | $3.1\ 10^{6}$ | $3.3\ 10^{4}$ | $4.4\ 10^{3}$ |
| $E_K^D$ (kJ $mol^{-1}$) | 26.2 | 23.8 | 21.4 | 9.7 | 5.2 |

**Supplementary Table 3: Arrhenius parameters for the five biomes studied in German *et al*. (2012)[6].**
